# An atypical endomembrane localized CNL-type immune receptor with a conserved deletion in the N-terminal signaling domain functions in cell death and immunity

**DOI:** 10.1101/2023.09.04.556214

**Authors:** Sruthi Sunil, Simon Beeh, Eva Stöbbe, Kathrin Fischer, Franziska Wilhelm, Aron Meral, Celia Paris, Luisa Teasdale, Zhihao Jiang, Lisha Zhang, Emmanuel Aguilar Parras, Thorsten Nürnberger, Detlef Weigel, Rosa Lozano-Duran, Farid El Kasmi

## Abstract

Plants have evolved intracellular nucleotide-binding leucine rich repeat receptors (NLRs) to induce a superior immune response. Upon activation, coiled-coil (CC) domain containing NLRs (CNLs) oligomerize to form apparent cation channels that promote calcium influx and cell death induction, with the alpha-1 helix of the individual CC domains penetrating membranes. Some members of a monophyletic subclass of CNLs, the ancient and autonomous NLRs (ANLs), are characterized by putative N-myristoylation and S-acylation sites at the N-terminus of their CC_G10/GA_ domain, potentially mediating permanent membrane association. Whether these **P**otentially **M**embrane **L**ocalized NLRs (PMLs) mediate cell death upon activation in a similar way as reported for other CNLs has been unknown. We integrated phylogenetic, cell-biological, and functional studies to uncover the cell death function of an atypical but conserved Arabidopsis PML, PML5, which has a 113 amino acid deletion in its CC_G10/GA_ domain. Active PML5 oligomers localize in Golgi membranes and the tonoplast, changes vacuolar morphology, and induce cell death, with the short N-terminus being sufficient for cell death. Mutant analysis supports a potential key role of PMLs in plant immunity. Similar deletions as in Arabidopsis PML5 are found in several Brassicales paralogs, pointing to the evolutionary importance of this innovation. PML5 is thus a naturally occurring CNL variant with a minimal signaling ‘domain’ and its further study should help in understanding the functional importance of this minimal domain for NLR signaling.

## INTRODUCTION

Plant immunity is mediated by two types of immune receptors: pattern-recognition receptors (PRRs) and nucleotide-binding leucine-rich repeat receptors (NLRs) (Ngou et al., 2021). PRRs detect conserved molecular patterns (pathogen-/microbe-/danger-associated patterns – P/M/DAMPs) at the cell surface and transmit this information to the inside of the cell via phosphorylation cascades. The downstream responses include reactive oxygen production, transient Ca^2+^ fluxes, and transcriptional reprogramming (Bigeard et al., 2015; Jeworutzki et al., 2010). NLRs detect the activity of pathogen-derived effector proteins (also known as virulence proteins) after they have been delivered into the host cell. Effector-triggered and NLR-mediated immunity is mostly a potentiation of PRR-mediated immune outputs (Ngou et al., 2021; Yuan et al., 2021). However, NLR activation often also leads to a specialized type of programmed cell death, the hypersensitive response (HR), of the infected cell (Balint-Kurti, 2019; El Kasmi, 2021). NLRs can be classified into four major subclasses based on their N-terminal domains: Toll-like/interleukin 1 receptor (TIR) domain containing TNLs, coiled-coil (CC) domain containing CNLs, RPW8-like CC (CC_R_) domain containing RNLs, and the ancient and autonomous CC_G10/GA_-type NLRs, or ANLs (Kourelis, 2022; Lee et al., 2021).

Many TNLs and CNLs have been shown to directly or indirectly sense effectors, whereas the two RNL subfamilies, the ACTIVATED DISEASE RESISTANCE 1 proteins (ADR1s) and N REQUIREMENT GENE 1 proteins (NRG1s), also referred to as helper NLRs, are required downstream of all TNLs and some sensor CNLs (Castel et al., 2019; Jubic et al., 2019; Saile et al., 2020; Wu et al., 2019). Helper NLR- independent CNLs can autonomously induce immune responses (Maruta et al., 2022). An example is the *Arabidopsis thaliana* (hereafter Arabidopsis) CNL HopZ-ACTIVATED RESISTANCE 1 (ZAR1), which confers resistance to a variety of bacterial pathogens (Baudin et al., 2017). Effector recognition induces conformational changes that allow AtZAR1 monomers to form a pentameric resistosome complex (Wang et al., 2019b; Wang et al., 2019a). The AtZAR1 resistosome functions at the plasma membrane (PM) where the alpha-1 helices at the extreme N-terminus of each of the five AtZAR1 CC domains penetrate the PM to form a cation-permeable channel important for Ca^2+^ influx. Calcium influx is in turn a prerequisite for AtZAR1-induced disease resistance and cell death (Bi et al., 2021; Wang et al., 2019b). A similar cell-death inducing mechanism has been postulated for the Arabidopsis RNLs AtADR1, AtNRG1.1 and AtNRG1.2, which in their (auto-)activated state also form higher-order oligomeric Ca^2+^ permeable channels at the plasma membrane (Feehan et al., 2023; Jacob et al., 2021; Wang et al., 2023). The importance of the CC, CC_R_ and CC_G10/GA_ domains for cell death activity is further supported by the observation that overexpression of these domains alone is often sufficient to induce cell death or autoimmune symptoms (Bolus et al., 2020; Casey et al., 2016; Jacob et al., 2018; Kim et al., 2018; Lee et al., 2022; Sarah M. Collier, 2011; Wang and Balint-Kurti, 2015; Wroblewski et al., 2018). Indeed, the first 29 amino acids of the *Nicotiana benthamiana* CNL NLR-REQUIRED FOR CELL DEATH 4 (NRC4), which make up the potentially membrane-penetrating alpha-1 helix, are sufficient to induce weak cell death when fused to YFP and transiently overexpressed (Adachi et al., 2019). Furthermore, exchanging the very N-terminal 17 residues of NRC4, corresponding to approximately half of the alpha-1 helix, with the corresponding region of several other unrelated CNLs revealed a conserved function of this region for CNL cell-death activity. This N-terminal region also contains conserved (mostly hydrophobic) residues that are essential for cell death and resistance function of AtZAR1 and NbNRC4 (Adachi et al., 2019).

Oligomerization and resistosomes formation have been demonstrated for several cell-death inducing CNLs and RNLs (Ahn et al., 2023; Bolus et al., 2020; Feehan et al., 2023; Forderer et al., 2022; Jacob et al., 2021; Li et al., 2020; Wroblewski et al., 2018), thus leading to the hypothesis that the CC/CC_R_/CC_G10/GA_ domains of all CNLs, including RNLs and ANLs, can form membrane-penetrating channels that promote cell death and immunity.

The ANL subclass of CNLs is characterized by the lack or severe degeneration of conserved motifs in the CC domain that have been implicated in cell-death activity of other CNLs (Lee et al., 2021). All ANLs that have been studied in details appear to be (plasma-)membrane localized and their CC_G10/GA_ domains alone can induce a cell death response when overexpressed (Kourelis et al., 2021; Lee et al., 2021; Wroblewski et al., 2018). The Arabidopsis Col-0 reference genome encodes 23 ANLs (Supplemental Table 1), including the well-characterized plasma membrane-localized proteins RESISTANCE TO PSEUDOMONAS SYRINGAE 2 (RPS2) and 5 (RPS5), SUPPRESSOR OF TOPP4-1 (SUT1), SUPPRESSOR OF MKK1 MKK2 2 (SUMM2), and the recently described L5 (Huang et al., 2021; Kourelis et al., 2021; Lee et al., 2021; Seo et al., 2016; Yan et al., 2019; Zhang et al., 2012).

Plasma membrane localization is important for CNL- and RNL-dependent immunity and cell death (Axtell and Staskawicz, 2003; Chen et al., 2017; El Kasmi et al., 2017; Qi et al., 2012; Saile et al., 2021; Takemoto et al., 2012). Seventeen of the 23 Arabidopsis ANLs feature residues subject to potential co- and/or post-translational modifications (PTMs) important for their (constitutive) (plasma-)membrane localization (Supplemental Table 1). For example, AtRPS5 is potentially N-myristoylated and S-acylated (S-palmitoylated) at glycine residues 2 and 3, as well as cysteine 4 (Qi et al., 2012). N-myristoylation is supposed to be an irreversible modification in which myristic acid is added to the N-terminus of proteins starting with methionine (M)–glycine (G)- X-X-X-serine (S)/threonine (T) residues, while S-acylation is a reversible modification usually involving the addition of a palmitate group to any cysteine (C) or serine(S) residue throughout the protein (Wang et al., 2021). S-acylation is thought to regulate membrane localization, protein-protein interactions, and cellular signaling processes (Hurst et al., 2023; Peitzsch and McLaughlin, 1993; Resh, 1999; Wang et al., 2021). AtRPS5 plasma membrane localization and cell death activity are lost when residues G2, G3 and C4 are mutated, consistent with a central role of PTMs at these residues (Qi et al., 2012). The extensive conservation of these residues and their importance for cell-death activity of ANLs raises an important question regarding the resistosome/cation channel model. How can these NLRs, or more specifically their CC_G10/GA_ domains, trigger cell death and immunity when their very N-terminus, preceding the alpha-1 helix, is permanently associated with the inner leaflet of the plasma membrane?

We have identified an ANL with a deletion of 113 amino acids at the N-terminus and potentially N-myristoylated glycine G2 and S-acylated cysteines C3 and C4 in the CC_G10/GA_ domain, which we named PML5 (**P**otentially **M**embrane **L**ocalized 5). PML5 provides a platform to study whether and how such variant ANLs function in cell death initiation and immunity. We demonstrate that PML5 is a canonical, self-associating ANL that forms activation-dependent higher-order oligomers. Overexpression of PML5 causes strong cell death in either *N. benthamiana* or Arabidopsis. Localization of (active) PML5 to Golgi membranes and the tonoplast, but not the cytosol, is required for cell death, which can be induced when only the first 60 amino acids of PML5 (CC^1- 60^) are overexpressed. PML5-like N-terminal deletions are also found in ANLs of other Arabidopsis accessions and in several Brassicales. Together, our results suggest that potentially constitutively membrane-associated ANLs have a canonical cell death function that requires their predicted alpha-1 helix and that the PML5-like ANLs may fulfil an important, evolutionary conserved function.

## RESULTS

### PML5 is an ANL with a conserved 113 amino acid deletion in its CC_G10/GA_ domain

N-terminal CC domains that form plasma membrane-penetrating cation channels are essential for immune signaling and cell death initiation (Adachi et al., 2019; Bi et al., 2021; Bolus et al., 2020; Forderer et al., 2022; Qi et al., 2012; Wang et al., 2019b; Wang et al., 2019a). We identified 20 CNLs in Arabidopsis Col-0 with putative N-myristoylation and/or S-acylation sites at their very N-terminus (Supplemental Table 1 and Figure 1A), which we named **P**otentially **M**embrane **L**ocalized (PML) NLRs. Seventeen of these PMLs belong to the monophyletic group of autonomous CNLs with a distinct CC_G10/GA_ domain, termed ANLs (Supplemental Figure S1 and Supplemental Table 1) (Kourelis et al., 2021; Lee et al., 2021). A detailed analysis of their coding sequences revealed that one of them, *PML5/At1g61300*, encodes for a 762 amino acid protein with a 113 amino acid deletion in its CC_G10/GA_ domain (Supplemental Figure S2). PML5 has all canonical motifs in its nucleotide-binding (NB-ARC) domain (Figure 1B) and forms a small subclade (cluster #3) with PML11, 12, and 13, which are all full-length PMLs (Supplemental Figure S1). AlphaFold 2 modelling of PML5 predicted a well-structured NB-ARC and C-terminal LRR domain, while the CC domain was predicted with low confidence as a short unstructured region followed by a single alpha-helix (amino acid 13-29) (Supplemental Figure 3A).

**Figure 1:**
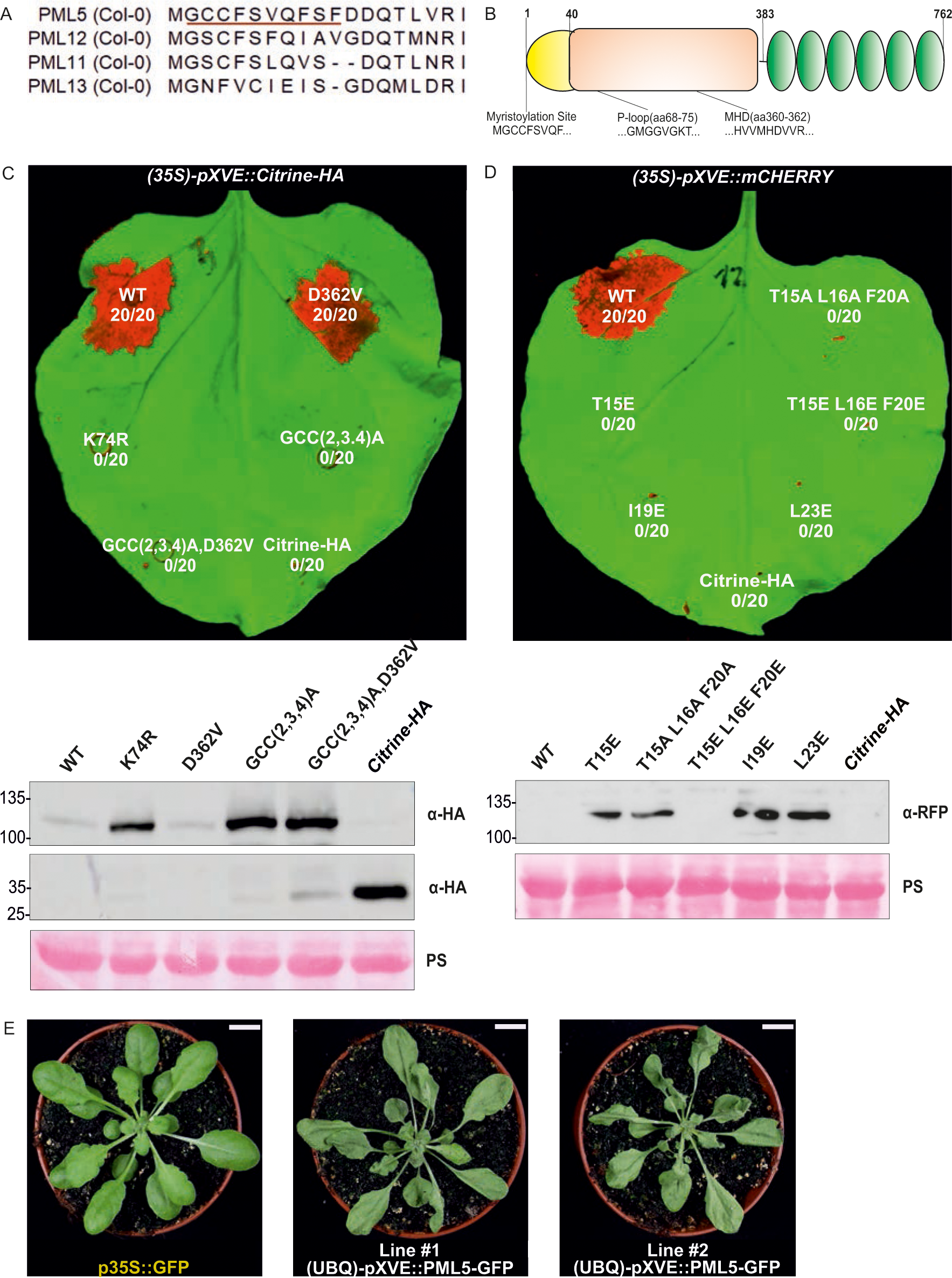
PML5 functions as a canonical cell death inducing CNL. (A) Sequence alignment of the first 23 N-terminal amino acids of PML5 related NLRs, highlighting the potential S-acylation and N-myristoylation sites. (B) Diagram of PML5 and its functional motifs. (C) and (D) Cell death induced by transiently expressed wild-type and mutant PML5 variants (top) in *N. benthamiana* leaves and corresponding protein blots (bottom, anti-HA (α-HA) antibody (C) and anti-RFP (a-RFP) antibody (D)), three days after induction with 20 µM estradiol. Citrine-HA was used as negative control. Bottom blot in (C) shows citrine-HA (about 30 kDa). Ponceau S (PS) staining shown as loading control. In (C), WT = wild-type PML5; D362V = MHD mutant; K74R = P-loop mutant; GCC(2,3,4)A = N-myristoylation and S-acylation (PTM) mutants; GCC/2,3,4)A,D362V = PTM / MHD double mutant. (E) Cell death in two independent transgenic Arabidopsis lines overexpressing *PML5-GFP*. A *35::GFP* plant was used as negative control. 4-week-old plants 30 hours after induction with 20 µM estradiol are shown. Scale bars, 1 cm.

We investigated the genomes of other Arabidopsis accessions (Jiao and Schneeberger, 2020; Liu et al., 2021; Van de Weyer et al., 2019) to determine whether other accessions also contain genes with a *PML5*-like deletion. In the genomes of 14 accessions, including Col-0 (at6909/Col-0; at9837/Con-0; at9721/Schip-1; at7413/Wil- 2; at9762/Etna-2; at1925/Che-2; at7416/Yo-0; at7058/Bur-0; at9814/Fell1-10; at9776/Fell3-7; at8285/Dralll-1; at6898/An-1; at7207/Kyoto; at9583/IP-Sne-0), a gene with high similarity to Col-0 *PML5*, including a similar deletion, was identified (Supplemental Figure S4). We also screened other plants for homologs with similar features, identifying several NLR proteins with a similar deletion and a potential N-myristoylation and S-acylation site in *Brassica napus* (BnaCnng57760D), *Capsella rubella* (XP_006300767.1; CARUB_v10019842mg; Carub.0002s0399), *Capsella bursa-pastoris* (Cbp33710), *Cardamine hirsuta* (CARHR052110), *Leavenworthia alabamica* (LA_scaffold903_5), and *Cleome vicosa* (UPZX_scaffold_2005671). A phylogenetic analysis suggested that the deletion in the CC domain may have arisen independently at least twice (Supplemental Figure S5). RNA-seq data indicate that *PML5* is expressed throughout the plant with the lowest expression in roots and the highest expression in rosette leaves (Klepikova et al., 2016; Zhang et al., 2020). These results demonstrate that ANLs with an N-terminal deletion and potential N-myristoylation and S-acylation sites are found throughout the Brassicales, suggesting a conserved and maybe important function.

### PML5 functions as a canonical cell death inducing CNL

Effector-dependent activation of CNLs often causes strong cell death (Chen et al., 2022; Chung et al., 2011; Li et al., 2019; Saur et al., 2019; Shao et al., 2003). NLR activation can be mimicked by introducing a single amino acid mutation in the conserved MHD motif of the NB-ARC domain, or in some cases by strong over- expression of wild-type NLRs (Adachi et al., 2023; Gao et al., 2011). To investigate whether PML5 can function as a cell-death inducing CNL, we transiently over- expressed PML5 wild-type and MHD mutant (D362V) proteins in *N. benthamiana*. PML5 and the D362V mutant elicited strong cell death regardless of the promoter or the C-terminal tag used (Figure 1C) (Supplemental Figure 3C and 3D). Mutating the conserved lysine at position 74 to arginine (K74R) in PML5 is expected to inhibit the P-loop motif function, which is essential for nucleotide binding and thus NLR activity (El Kasmi et al., 2017; Tameling et al., 2002; Williams et al., 2011). Indeed, the introduction of the K74R mutation abolished cell death (Figure 1B and 1C, Supplemental Figure 3C and 3D). This was not due to lesser accumulation of the K74R mutant (Figure 1C, Supplemental Figure 3C and 3D). Together, these analyses demonstrated that PML5 functions as a canonical CNL, although it lacks most of the CC domain residues found in other CNLs.

The N-terminal myristoylation and S-acylation sites of the well-studied PML RPS5/PML7 are important for RPS5/PML7 immune functions (Qi et al., 2012). We wanted to learn whether mutation of the predicted N-myristoylation and S-acylation sites of PML5 also affects its ability to cause cell death, and found that the GCC(2,3,4)A and GCC(2,3,4)A,D362V mutants did not induce any cell death, in spite of being well expressed (Figure 1C and Supplemental Figure 3C and 3D), suggesting that N-myristoylation and S-acylation are required for PML5 cell-death activity or localization. Conserved hydrophobic residues in the alpha-1 helix of the CC domain have recently been shown to be important for the cell-death activity of many CNLs (Adachi et al., 2019; Wang et al., 2019). Amino acids 13 to 29 of the PML5 N-terminus are predicted to form an alpha helix that aligns with the amino acids of the alpha-1 helix of Arabidopsis ZAR1 and *N. benthamiana* NRC4 (Supplemental Figure 3A and 3B). The alignment suggested that PML5 shares conserved hydrophobic residues in similar positions. To test whether the putative alpha-helix of PML5 is also contributing to PML5 cell death activity, we mutated several residues in this region (T15, L16, I19, F20 and L23), including the conserved hydrophobic ones (Supplemental Figure 3B). As shown for AtZAR1 and NbNRC4, mutating these residues individually or in combinations of three to alanine or glutamate completely abolished PML5 cell death activity (Figure 1D) (Adachi et al., 2019; Wang et al., 2019b). Protein blots confirmed the expression of each single mutant and the T15A L16A F20A triple mutant. The T15E L16E F20E mutant, however, could not be detected, possibly due to instability. We conclude that like other PMLs, PML5 cell death activity requires these conserved (hydrophobic) residues in the N-terminal putative alpha-1 helix.

To determine if PML5 cell-death activity is P-loop-dependent in Arabidopsis, as it is in *N. benthamiana*, we generated stably transformed Col-0 plants with *UBIQUITIN10* promoter-driven estradiol-inducible *PML5-GFP* wild-type, MHD- and P-loop mutant constructs. Induced expression of wild-type PML5 and the D362V mutant, but not the K74R P-loop mutant, caused severe cell death in leaves of four-week-old plants (Figure 1E and Supplemental Figure 3E). Additionally, seedlings expressing wild-type PML5 or the D362V mutant were severely stunted and eventually died when germinated and grown on estradiol-containing medium (Supplemental Figure 3F).

Together these data suggest that PML5, despite the 113 amino acid deletion in its CC domain, can function as a canonical CNL to induce cell death when activated or transcriptionally induced. PML5 is thus a naturally occurring CNL with a minimal signaling domain and therefore makes for an excellent platform to understand the functional importance of this minimal domain, including its structure, for NLR signaling.

### EDS1 and RNLs are not required for PML5 cell death activity

Many NLRs, including some effector sensing CNLs, require RNLs and the plant-specific lipase like proteins EDS1 (ENHANCED DISEASE SUSCEPTIBILITY 1), PAD4 (PHYTOALEXIN DEFICIENT 4) and SAG101 (SENESCENCE ASSOCIATED GENE 101) to induce cell death and immunity (Castel et al., 2019; Jubic et al., 2019; Lapin et al., 2020; Saile et al., 2020; Sun et al., 2021; Wu et al., 2019) To investigate whether PML5 also requires RNL or EDS1 function, we transiently expressed wild-type PML5 and the D362V mutant in *eds1*, *adr1*, *nrg1* single and *adr1 nrg1* double mutant *N. benthamiana* plants. In all mutant backgrounds, both wild-type PML5 and its MHD mutant induced cell death comparable to the cell death in wild-type *N. benthamiana* leaves (Supplemental Figure 6), indicating that PML5 cell death activity is independent of RNL helper NLRs and EDS1.

### The N-terminal 60 amino acids of PML5 are sufficient for cell death

Overexpression of the full-length CC domain of many CNLs is sufficient to induce strong cell death, demonstrating that the N-terminal CC domain is the signaling, potentially pore-forming domain (Adachi et al., 2019; Baudin et al., 2020; Chia et al., 2022; Li Wan, 2019; Maekawa et al., 2011; Wang et al., 2020). To determine whether the N-terminal partial CC domain of PML5 is sufficient for cell death we generated different PML5 truncations (Figure 2A). Based on the predicted AlphaFold 2 structure, we constructed three different CC-type fragments (CC^1-35^, CC^1-39^ and CC^1-60^), three NB-ARC containing fragments (NB^36-363^, NB^40-383^ and NB^61-383^) and two LRR fragments (LRR^364-762^ and LRR^384-762^) and expressed these variants in *N. benthamiana*. None of the NB-ARC or LRR containing fragments were able to cause visible cell death (Figure 2B and Supplemental Figure 7A), but PML5 CC^1-60^ induced cell death, albeit weakly (Figure 2B and Supplemental Figure 7A). Lack of cell death was not due to lack of expression, as confirmed with protein blots (Figure 2B (bottom panel)).

**Figure 2:**
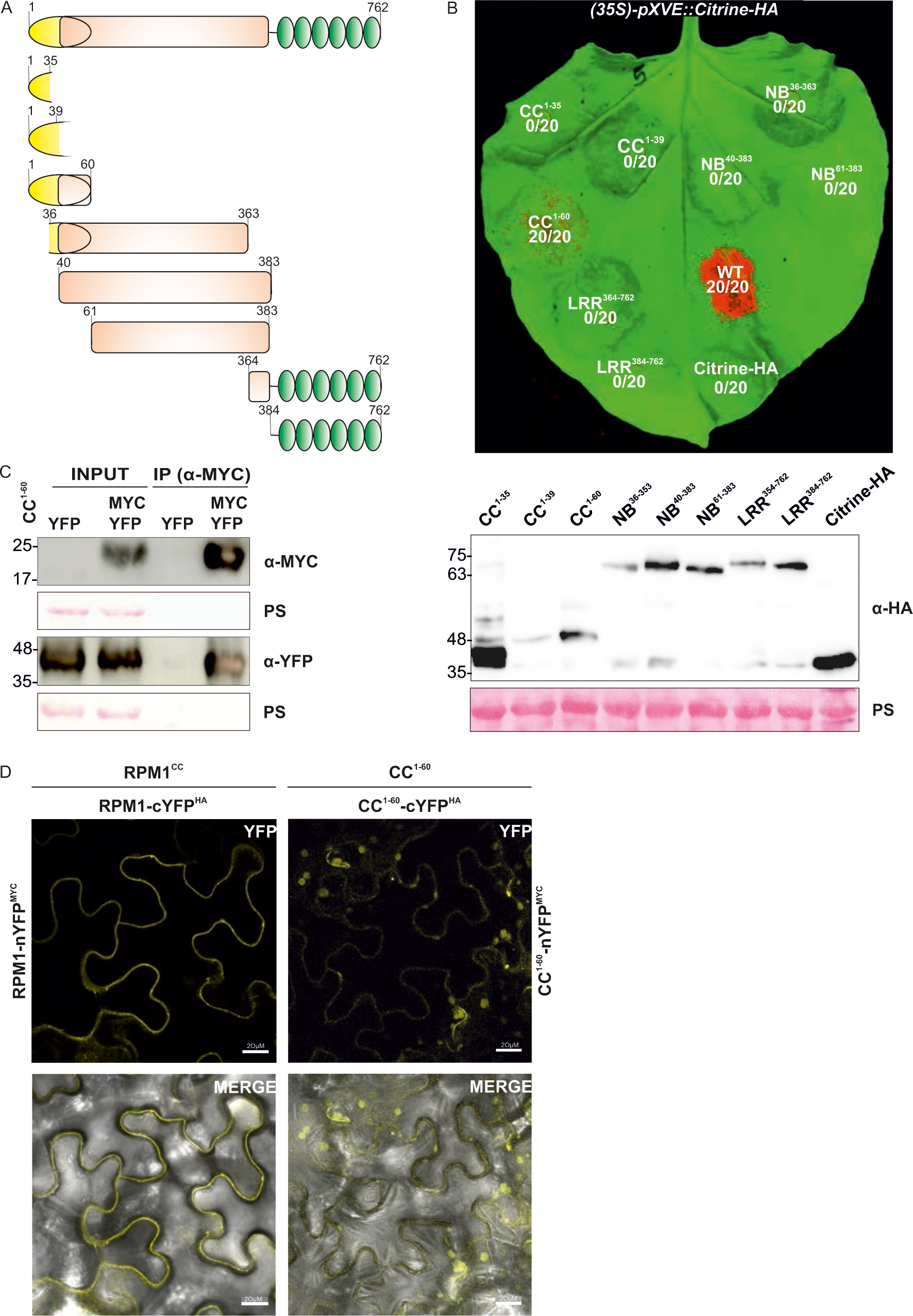
PML5 N-terminal 60 amino acids self-associate and are sufficient for cell death induction. (A) Diagram of full-length PML5 and its individual domains. (B) Cell death induced by transiently expressed full-length PML5 and PML5 domains in *N. benthamiana* (top). Citrine-HA was used as negative control. Photos were taken two days after induction with 20 µM estradiol. Fusion proteins were detected with an anti-HA (α-HA) antibody. Ponceau S (PS) staining shown as loading control. (C) PML5 CC^1-60^ self-associates, as shown by transient expression in *N. benthamiana*. C-terminal Myc- and YFP-CC^1-60^ fusions were co-expressed. Proteins were immunoprecipitated using anti-myc (α- MYC) beads and detected using anti-myc (α-MYC) and anti-GFP (α-GFP) antibodies. Ponceau S (PS) staining shown as loading control. (D) Bimolecular fluorescence complementation (BiFC) of PML5 CC^1-60^ fusion proteins, CC^1-60^-cYFP^HA^ and CC^1-60^- nYFP^MYC^ transiently expressed in *N. benthamiana*. RPM1 CC^1-155^ served as positive control. Scale bars, 20 µm.

**Figure 3:**
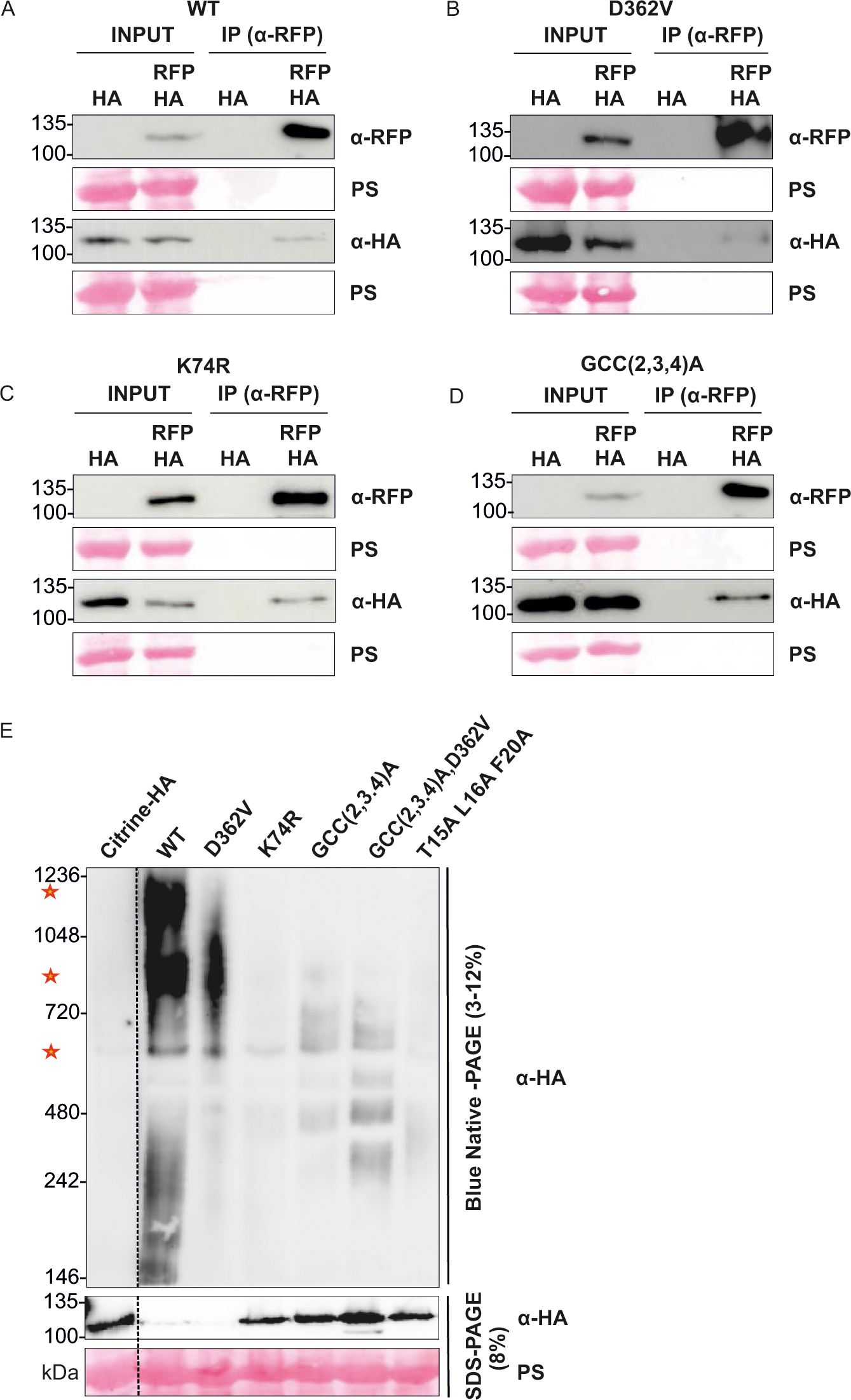
Full-length PML5 self-associates and forms high-molecular weight complexes. Full-Length PML5 (A), D362V MHD mutant (B), K74R P-Loop mutant (C), GCC(2,3,4)A) (D) self-associate, as seen after transient expression in *N. benthamiana*. PML5-citrine-HA and PML5-mCherry expression was induced with 20 µM estradiol. Proteins were extracted four hours after induction and immunoprecipitated using anti-RFP (α-RFP) beads and detected using anti-RFP (α- RFP) and anti-HA (α-HA) antibodies. Ponceau S (PS) staining shown as loading control. (C) PML5 wild-type and D362V mutant form high-molecular weight complexes. Total protein from plants transiently expressing full-length C-terminally tagged citrine- HA PML5 and PML5 mutants was extracted 6 hours after induction with 20 µM estradiol, separated by BN-PAGE, and detected with an anti-HA (α-HA) antibody. Protein extracts were also separated by SDS-PAGE and protein detected with anti- HA (α-HA) antibody. Ponceau S (PS) staining shown as loading control. Red stars indicate potential PML5-containing complexes of different sizes.

We wanted to test whether different PML5 domains could complement each other in trans, as described for the *Solanaceae* CNL Rx (Peter Moffett, 2002), and co-expressed PML5 CC^1-60^ with the corresponding NB-ARC^61-383^ and LRR^384-762^ fragments. Full cell death activity could not be achieved (Supplemental Figure 7B), indicating that the first 60 amino acids of PML5 are capable of cell death induction, but not to the same level as the full-length protein.

Activation of and thus cell death induction by CNLs or their CC domains requires self-association or oligomerization (El Kasmi et al., 2017; Wang et al., 2019b). We aimed to determine whether the cell death inducing PML5 CC^1-60^ fragment is also self-associating. Co-immunoprecipitation (Co-IP) and bimolecular fluorescence complementation (BiFC) experiments of transiently expressed CC^1-60^ revealed self- association of the 60 amino acids fragment (Figure 2C and 2D) (Supplemental Figure 7C and 7D). Thus, the PML5 CC domain with the 113 amino acid deletion is sufficient for cell death induction.

### PML5 constitutively self-associates and forms higher-order complexes upon activation

Next, we sought to investigate whether the full-length wild-type and D362V mutant PML5 proteins self-associate. Co-IP experiments of differentially tagged PML5 indicated that not only the wild-type and D362V proteins self-associate (Figure 3A and 3B), but also the non-functional K74R and GCC(2,3,4)AAA mutants (Figure 3C and 3D). Attempts to support the co-IP results by BIFC analysis failed, even though the tagged proteins were all expressed (data not shown). This may be due to steric hindrance of the fluorophore complementation when the PML5 proteins are tagged C-terminally.

Association of two proteins in co-IP experiments suggests close spatial proximity of these proteins but does not necessarily demonstrate their ability to form oligomers, as hypothesized for activated CNLs. Blue native PAGE (BN-PAGE) is a suitable approach to examine the complex formation and thus the oligomerization of proteins and has already been used to successfully show CNL resistosome formation upon (auto-)activation (Ahn et al., 2023; Feehan et al., 2023; Hu et al., 2020; Jacob et al., 2021). To test if PML5 forms resistosome-like complexes in an activation dependent manner we performed BN-PAGE with transiently expressed PML5 wild-type and mutants (Figure 3E). The detection of protein complexes at ∼550-600 kDa for all PML5 protein variants confirmed the co-IP results, suggesting that regardless of activation status some PML5 proteins may form or be part of such complexes. Above 720 kDa and between 1048 kDa and 1236 kDa, we detected the presence of two PML5 containing complexes, especially for PML5 wild type and D362V. The potential PML5 complex above 720 kDa was also detected very faintly with the GCC(2,3,4)A and GCC(2,3,4)A,D362V mutants (Figure 3E), indicating that the potential PTM sites at the N-terminus are not required for oligomerization of PML5. We did not detect the formation of the two complexes above 720 kDa for the K74R and the T15,L16,F20A loss-of-function mutants, suggesting requirement of both a functional P-loop and the conserved residues in the potential alpha-1 helix for oligomerization, but not for self-association. Altogether, co-IP and BN-PAGE results indicated that only active PML5 forms a resistosome-like complex or at least associates with other proteins to form a high-molecular weight complex that can induce cell death.

### PML5 localizes to Golgi’s and the tonoplast in a dynamic manner

Classification of PML NLRs based on their N-terminal myristoylation and/or S-acylation site(s) (Figure 1A and Supplemental Table 1) suggested that they are PM localized. In fact, all PMLs characterized thus far have been reported to be PM associated (Huang et al., 2021; Qi et al., 2012; Yan et al., 2019; Zhang et al., 2017). Confocal laser scanning microscopy of conditionally expressed PML5-Citrine-HA in *N. benthamiana* leaves indicated that PML5 wildtype exhibits a dynamic localization. We observed PML5 on Golgi membranes six hours post induction (hpi) (Figure 4A) and at the tonoplast (Figure 4B), Golgi and cytosol 7 hpi and later (Figure 4C). We could not detect any localization at the plasma membrane at the time points analyzed (Figure 4D), which are just before cell death became macroscopically and microscopically visible. We confirmed the membrane localization of wild-type PML5 by subcellular fractionation experiments (Supplemental Figure 8E). At the timepoint PML5 was localizing to tonoplast membranes, we observed a dramatic change in vacuolar morphology (Figure 4B and 4C). All PML5 expressing cells displayed this change in vacuolar morphology, a vesiculation or vacuolar fragmentation that was severely enhanced when the PML5 D362V mutant was expressed (Supplemental Figure 8D). These findings indicate a strong correlation of this vacuolar phenotype with the cell death activity of PML5. Stable transgenic Arabidopsis lines conditionally expressing PML5-GFP showed a similar vacuolar fragmentation phenotype in elongating primary root cells (Figure 4E). The PML5 loss-of-function mutants K74R, GCC(2,3,4)A and GCC(2,3,4)A,D362V localized to the cytosol with no detectable membrane localization, and did not induce any vacuolar fragmentation phenotype (Supplemental Figure 8A,8B and 8C).

**Figure 4:**
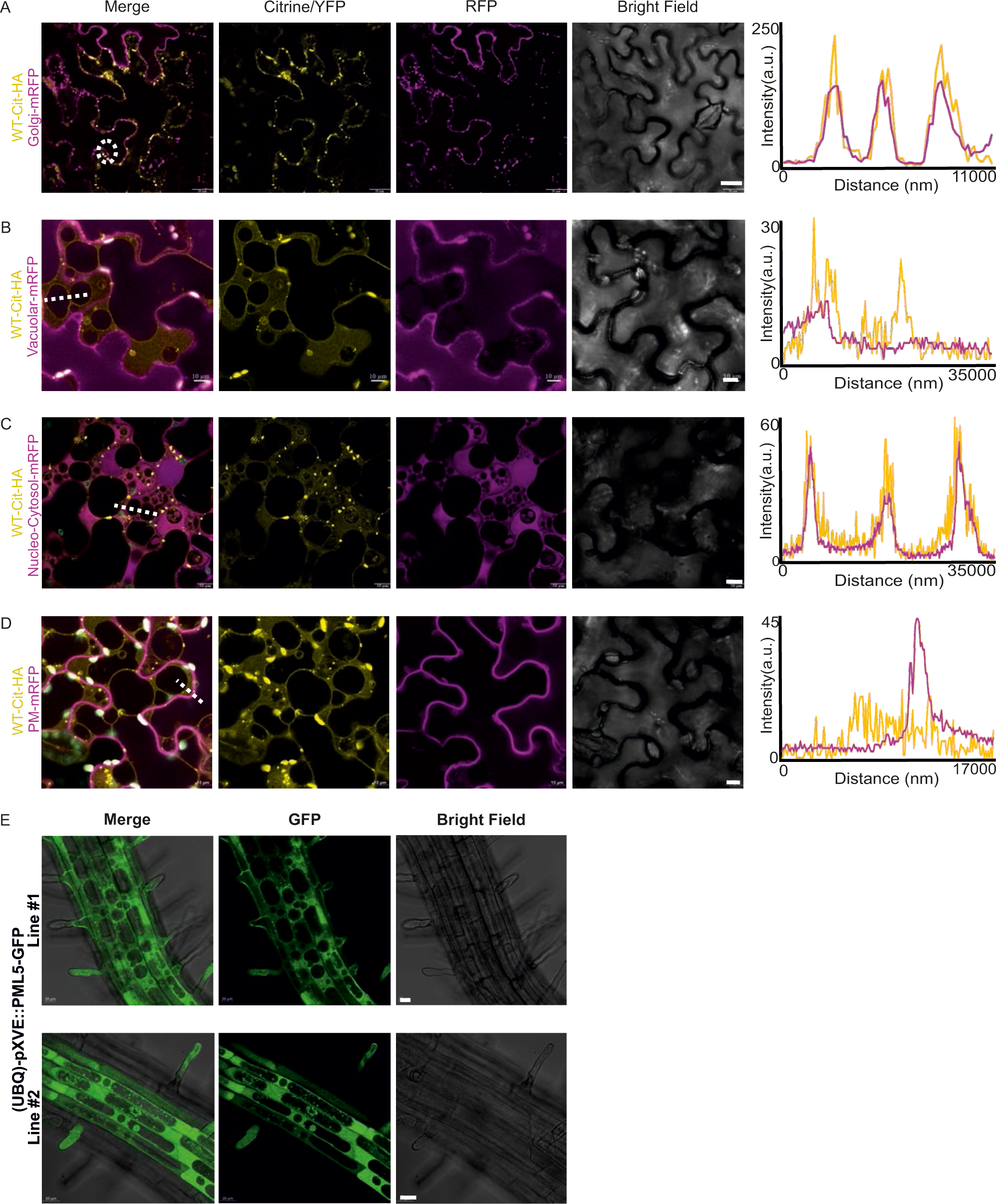
PML5 localizes to different cellular compartments. (A-D) Confocal laser scanning microscopy indicates localization of transiently expressed PML5-citrine-HA to Golgi membranes (A), the vacuolar membrane (B), and cytosol (C), but not to the plasma membrane (D). White dotted lines indicate the area used for colocalization profile analysis and the corresponding profiles are shown on the right, except for (A), where a line crossing the three circled puncta was used to extract the intensity profile. Scale bars, 10 µm; see Methods for compartment marker proteins. PML5-citrine-HA expression was induced with 20 µM estradiol and confocal microscopy was performed 6-8 hours after induction: Golgi (6 hpi), vacuolar membrane (7 hpi), cytosol (7 hpi), plasma membrane (8 hpi). (E) PML5-GFP localization in Arabidopsis elongating root cells. Scale bars, 20 µm. PML5-GFP expression was induced with 20 µM estradiol and confocal microscopy was performed 18 hours after induction.

Our analyses suggest a dynamic localization of PML5, being associated with Golgi membranes first and then detectable at the tonoplast and cytosol. Golgi and or tonoplast localization seems to be required for cell death activity, which is preceded by vacuolar fragmentation. Whether this vacuolar fragmentation is the cause or consequence of PML5-induced cell death is not yet clear.

### Loss of *PML5* in Col-0 modestly increases susceptibility to bacterial infection

As reported above, induced (over-)expression of active/activated PML5 strongly induced cell death and altered vacuolar morphology (Figure 1 and 4), suggesting that PML5 functions as a canonical CNL but with some specific characteristics - a dynamic Golgi/tonoplast localization and the vacuolar fragmentation phenotype. To elucidate whether loss of *PML5* would have any effect on development or immunity, we used CRISPR/Cas9 gene editing to generate a knockout mutant, *pml5-2c*, with a 4,192 bp deletion, spanning the entire coding sequence, which is replaced by a short 6 bp stuffer fragment (Figure 5A and see material and method section). We characterized this mutant together with a *pml5* T-DNA insertion line, *pml5-1* (Salk_0691926) (Figure 5A). The *pml5* mutants were morphologically indistinguishable from Col-0 wild type (Figure 5B and C) but flowered early in our green house and growth chambers (Figure 5D). The *pml5-1* allele is not a full knockout, with some remaining transcripts generated from sequences upstream of the T-DNA insertion (Figure 5E).

**Figure 5:**
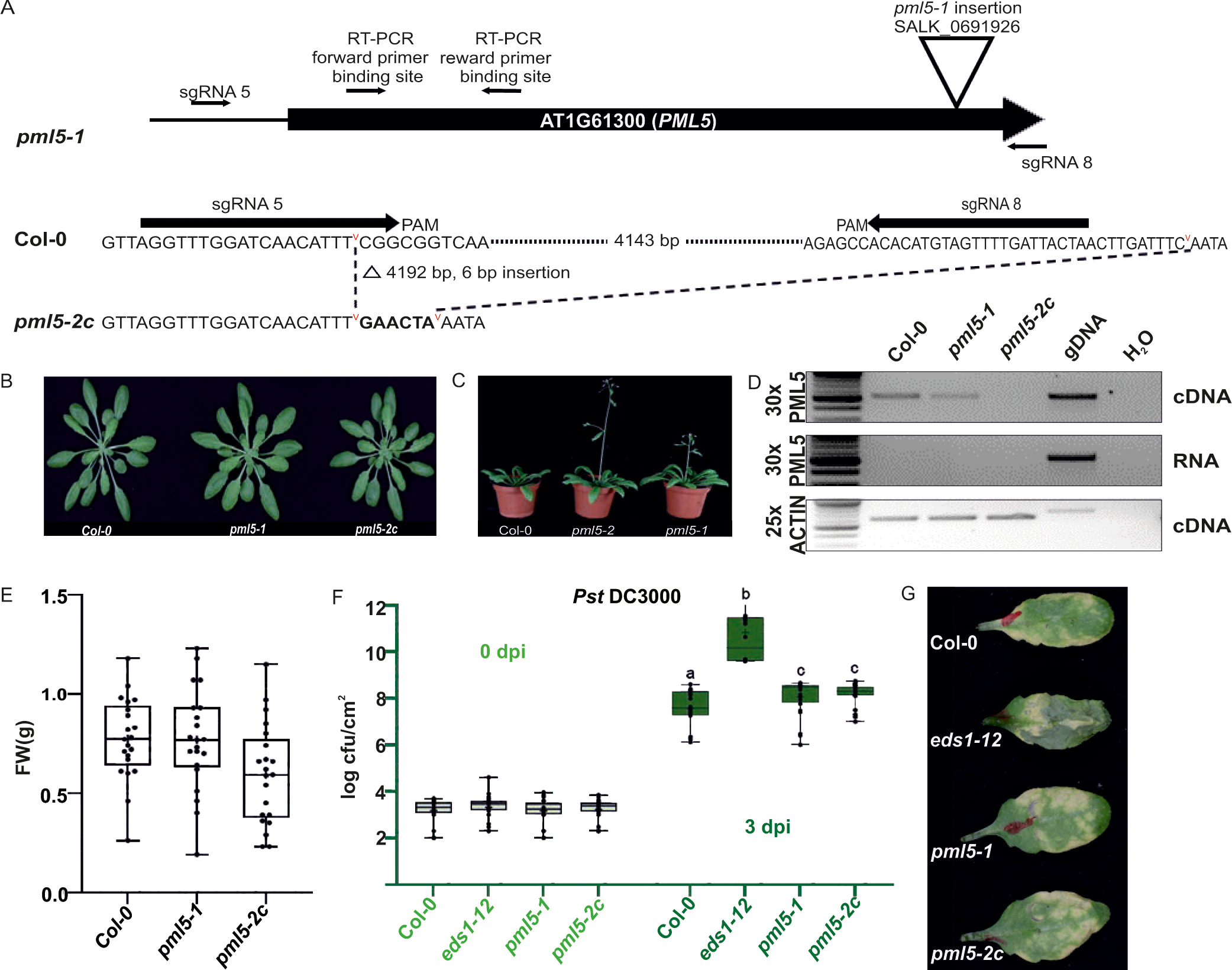
*pml5* mutants are more susceptible to *Pst* DC3000 infection. (A) Diagram of *pml5* alleles. SALK T-DNA insertion (*pml5-1; SALK_0691926*) and CRISPR/Cas9 induced 4,192bp deletion/6bp insertion in *pml5-2c*. (B) Rosette of eight- week-old Arabidopsis *pml5-1* and *pml5-2c* mutants compared to wildtype Col-0 grown in short days. (C) Fresh weight of eight-week-old wild-type*, pml5-1,* and *pml5-2c* (*n*=21) plants from panel (B). There was no significant difference (Tukey’s HSD test). (D) Early flowering of four-week-old *pml5-1* and *pml5-2c* mutants. Plants were grown in long days. (E) No detectable *PML5* transcripts in *pml5-2c* and reduced transcript levels in *pml5-1*, as determined by RT-PCR (30 cycles). *ACTIN2* (25 cycles) is used as positive control and gDNA (30 cycles) as negative control. (F) Six-week-old plants, hand infiltrated with *Pst* DC3000 (OD_600_ = 0.001) support modestly enhanced bacterial growth in *pml5-1* and *pml5-2c*. Infiltrated leaves were collected on day 0 and day 3 for bacterial colony counts. Data points (colony forming units per square cm – cfu/cm^2^) represent six biological replicates with four technical replicates each (*n*=24, each with six leaf samples). Significance from Tukey’s HSD (α = 0.05). (G) *Pst* DC3000-induced disease symptoms on *pml5-1* and *pml5-2c*.

To learn whether *PML5* has any function during immunity induced upon infections with a bacterial pathogen, we infected them with the virulent *Pseudomonas syringae* pv. *tomato (Pst)* DC3000 strain. Both *pml5* alleles were slightly more susceptible to *Pst* DC3000 infections (Figure 5F). This was also visible on the infected leaves, as disease symptoms were slightly more pronounced in *pml5-1* and *pml5-2c* compared to wild-type plants (Figure 5G).

To understand if *PML5* is involved in ETI responses or in basal immunity, we used *Pst* DC3000 expressing effectors that activate specific NLRs (RPM1, RPS5, ZAR1 and RPS4/RRS1) in Col-0 and the effector depleted *Pst DC3000* D36E strain (Debener et al., 1991; Lewis et al., 2010; Narusaka et al., 2009; Simonich and lnnes, 1995; Wei et al., 2015). Sensor NLR-induced bacterial growth restriction and HR was not altered in *pml5-1* and *pml5-2c* mutants (Supplemental Figure 9A – H). Similarly, we did not observe any effect on growth restriction of *Pst* D36E on *pml5* mutants (Supplemental Figure 9A – H). The lack of any measurable effect on *Pst* D36E growth in *pml5* indicates no important role for PML5 in PRR-mediated immunity. Indeed, reactive oxygen species (ROS) accumulation and ethylene production, two major PRR-induced responses, upon treatment of *pml5-2c* with the PAMPs flg22 (triggers the PRR FLS2) (Mersmann et al., 2010) or nlp20 (triggers the PRR RLP23) (Bohm et al., 2014) were not negatively affected (Supplemental Figure 10A and B).

The immune response against the virulent bacterial pathogen *Pst* DC3000 was marginally affected by loss of PML5 (Figure 5F). This could indicate a general function of PML5 for immunity that is not observable during immunity against avirulent or effector-less *Pst* DC3000 D36E. To investigate this hypothesis further, we inoculated both *pml5* mutants and Col-0 with tobacco rattle virus (TRV) (Liu et al., 2002). TRV accumulation, measured by quantification of viral RNA in the infected leaf tissue, was not significantly altered compared to wild type (Supplemental Figure 10C), indicating PML5 does not contribute to anti-TRV resistance. Our infection experiments with different PAMPs and pathogens suggests that PML5 has no specific function in basal immunity, the tested ETI responses or in resistance against TRV.

## DISCUSSION

We report the identification and initial characterization of a CC_G10/GA_-type NLR (ANL) immune receptor with a conserved deletion of most of the N-terminal CC signaling domain. Our analyses demonstrate that despite its 113 amino acid deletion, PML5 functions as a canonical full-length ANL in respect to the cell-death initiation mechanism. PML5 has strong, P-loop dependent cell-death activity when overexpressed (Figure 1) and residues 1 to 60 are sufficient for cell death induction, suggesting that the PML5 N-terminal region is the cell death executing part. This is surprising, because the PML5 N-terminal domain is most likely N-myristoylated and S-acylated, potentially causing constitutive membrane association of PML5. AlphaFold 2 structural prediction suggests that the PML5 CC_G10/GA_ domain also lacks three of the alpha-helices normally found in full-length CNLs and ANLs (Bentham et al., 2018; Forderer et al., 2022). However, several hydrophobic residues that were recently suggested to be required for proper function and oligomerization of all cell death inducing CNLs (Chia et al., 2022) are retained in the PML5 N-terminal signaling/executioner domain and required for PML5 function (Figure 1C). Thus, the remaining predicted alpha helix of PML5 may serve the same function as the alpha-1 helix of AtZAR1 or NbNRC4.

The potential N-myristoylation and S-acylation sites in PML5 are shared by full-length members of the PML family and are important for at least RPS5/PML7 (Qi et al., 2012) and L5 function (Huang et al., 2021). In our experiments, mutation of PML5 G2, C3 and C4 affected cell death activity and Golgi/tonoplast localization, suggesting that PML5 activity also requires this post-translational modification. We hypothesize that pre-activation localization at the correct cellular membrane, like the PM or other endomembrane compartments, is mediated by these modifications and is required for proper activation and therefore function of all PML NLRs. This is supported by our findings of lack of cell-death activity of the G,C,C(2,3,4)A,D346V mutant. Potentially, this PTMs are also enabling or at least supporting oligomerization of activated PML5 (Figure 3E). How such acylated NLRs are capable of inducing cell death by a similar mechanism as suggested for CNLs or RNLs is still not fully clear. It is possible that a de-palmitoylation of the modified cysteines or maybe a phosphorylation of residues close to the PTM sites could release the very N-terminus from the inner leaflet of the membrane, thus enabling it to oligomerize and penetrate the lipid bilayer for channel or pore formation upon activation. Cleavage of the very N-terminal region upon or before activation by an unknown protease may also be possible. However, we did not observe a size difference of cell death active wildtype or D364V mutant PML5 compared to the PML5 loss of function variants in our protein blots. The detection of such a very small N-terminal cleavage product would be difficult, since N-terminal tagging of CNLs does negatively affect their cell death function and in the case of PMLs their S-acylation/N-myristoylation and thus their localization, which is required for their activity. Our results and previous published work of other PMLs indicate that this family of CNLs most likely induces cell death by a very similar mechanism as CNLs with a so-called MADA, MADA-like or MAEPL motif at their very N-terminus (Adachi et al., 2019; Chia et al., 2022).

Oligomerization leads to pore or cation-permeable channel formation at the PM and Ca^2+^ influx, and these are required for cell death and immune function for several CNLs and the RNL protein family (Bi et al., 2021; Contreras et al., 2023; Forderer et al., 2022; Jacob et al., 2021; Wang et al., 2019b; Wang et al., 2019a). All PMLs characterized so far localize at the PM where they execute their cell death function (Huang et al., 2021; Qi et al., 2012; Yan et al., 2019; Zhang et al., 2017). In agreement, we could show that PML5 self-associates and that it oligomerizes upon activation (Figure 3), potentially forming resistosomes like other CNLs.

PML5 is, to our knowledge, the first (Arabidopsis) CNL found to be Golgi and tonoplast localized. Our results strongly suggest that PML5 executes its cell death function at one of these compartments, most likely at the tonoplast. PML5 could act as a cation or Ca^2+^ permeable channel at the tonoplast enabling the release of Ca^2+^ from the vacuole into the cytoplasm, The vacuole, like the endoplasmic reticulum and the apoplast, is an important storage of calcium in plants (Peiter, 2011; Schonknecht, 2013), and could source Ca^2+^ required for NLR-triggered cell death. The interpretation that PML5 may release Ca^2+^ from the vacuole into the cytoplasm is supported by the changed vacuolar morphology when cell-death active PML5 is overexpressed. This vacuolar fragmentation/fission or tubulation lead to the occurrence of bubble-like structures and in some cases also an obvious membrane invagination (Figure 4), which is very similar to what was recently been described for the overexpression of plant mechanosensors of the PIEZO protein family (Radin et al., 2021). Here, tonoplast localized PIEZOs promote the release of Ca^2+^ stored in vacuoles and modulate vacuolar complexity/morphology enabling the plant cell to respond to changes in cell mechanics. Whether there is a relation between PML5-induced changes in vacuolar morphology and plant PIEZOs remains to be elucidated.

How did the deletion of most of the CC domain arise independently at least twice during evolution and why were PML5 homologs with such deletions not lost during evolution? We currently can only speculate about potential answers. PML5 and its four closest orthologs (PML11, 12 and 13) belong to the class of highly variable NLRs (hvNLRs) in Arabidopsis (Prigozhin and Krasileva, 2021). hvNLRs are specified by high allelic diversity, especially in the LRR coding region. It is conceivable that the nucleotide sequence at the 5’ end (CC domain encoding part) of these PMLs is also prone to deletions because of the presence of fragile DNA sites, similar to recurrent deletions that confer evolutionary advantages that have been found in other systems (Xie et al., 2019). This would be consistent with the finding that gene body mutations in Arabidopsis are more likely at the 5’ and 3’ ends of genes (Monroe et al., 2022). hvNLRs have been hypothesized to act as sources for novel recognition specificities of NLRs (Prigozhin and Krasileva, 2021), and a specific effector (or pathogen) recognized by AtPML5 may not exist. In any case, the occurrence of PML5-like deletions in NLRs of other Arabidopsis accessions and Brassicales species indicates an evolutionary advantage of retaining this mutation (deletion) in the population of diversified species. PML5-like NLRs might serve an important function in the immune system and are under balancing selection (Koenig et al., 2019). Balancing selection to maintain intra- and interspecific allelic diversity was recently described for a barley NLR with a novel integrated Exo70 domain that was also found in NLRs of unrelated *Poaceae* species, which have diversified from barley 24 million years ago (Brabham et al., 2018). Balancing selection has also been suggested for certain paired NLRs (Shimizu et al., 2022).

The precise function of PML5 and PML5-like NLRs in immunity needs further investigation. We observed a weak decrease in resistance against the hemibiotrophic pathogen *Pst* DC3000 for *pml5* mutants under our growth conditions (Figure 5). This was already previously reported for *pml5* and mutants deficient for other PMLs in Arabidopsis, including *RPS5/PML7* (Jiang et al., 2020). The transcription of many PMLs is regulated by the microRNA472-mediated silencing pathway and overexpression or knockout of miR472 decreases or enhances basal disease resistance against *Pst* DC3000, respectively (Boccara et al., 2014; Jiang et al., 2020). PMLs may function in or contribute to basal resistance in general, or they may have evolved to recognize specific effectors or guard conserved effector targets (RPS5/PML7; SUMM2/PML8; SUT1/PML17).

## METHODS

### Transgenes

The coding sequences of *PML5*, *eYFP-HA,* and eYFP were cloned into GATEWAY^TM^ compatible vector pDONOR207 (Invitrogen, Carlsbad, USA) via GATEWAY^TM^ Cloning Technology (Thermo Fischer Scientific, Waltham USA) using primers listed in Table S1. All mutant constructs were generated by Site-Directed mutagenesis PCR using primers listed in Table S1. The PCR products were digested overnight with *DpnI* (NEB, Frankfurt am Main, Germany) and transformed into *Escherichia coli* DH5α. All constructs were confirmed by sanger sequencing (Eurofins, Ebersberg, Germany). LR reactions (Gateway Cloning^TM^ Technology Thermo Fischer Scientific, Waltham USA) were performed to introduce the CDS into a modified 35S promoter-driven estradiol-inducible destination vector pMDC7-Citrine-HA (Curtis and Grossniklaus, 2003), Ubiquitin promoter driven estradiol-inducible destination vector pMDC7-GFP, estradiol-inducible destination vector pABindmCherry (Bleckmann et al., 2010), 35S promoter driven destination vector pGWB641, pGWB617 (Nakamura et al., 2010) and pBAT-N and pBAT-C BIFC vectors. The different domain constructs for PML5 were amplified from full-length CDS with primers listed in Table S1. Gateway Cloning^TM^ was performed as described above.

### Plant material and growth conditions

*A. N. benthamiana* plants used in the study were grown in pots with soil for 4-5 weeks in a walk-in growth chamber under the following conditions: 12-hour light/12-hour dark, 24°C/22°C, with relative humidity (RH) of up to 70%. *N. benthamiana* mutant lines *eds1* (Schultink et al., 2017), *nrg1* (Qi et al., 2018), *adr1*, *nrg1, adr1 nrg1* were obtained from Jeff Dangl’s lab. *A. thaliana* plants were grown in walk-in growth chambers with short day (8-hour light/16-hour dark in 21°C/18°C, 45% RH) or long day conditions (16-hour light/8-hour dark in 21°C/18°C, 45% RH).

### Transgenic plants

*A. thaliana* Col-0 plants at the flowering stage were dipped with *Agrobacterium tumefaciens* strain GV3101/pMP90 carrying the specific constructs (Clough and Bent, 1998). Transformants were selected on ½ Murashige and Skoog (MS) agar plates containing 15ug/ml Hygromycin (Invitrogen, Thermo Fisher Scientific, Carlsbad, CA). Transgene expression was checked in T3 plants by immunoblotting total protein extracts with α-GFP antibodies.

### CRISPR/Cas9 genome editing

The sgRNA sequences unique to the *PML5* locus with predicted high efficiency were selected using CCTOP (Labuhn et al., 2018; Stemmer et al., 2015). Dicot genome editing vectors (pDGE) were used as the cloning system (Grutzner et al., 2021; Ordon et al., 2017; Stuttmann et al., 2021). Oligonucleotides containing the *PML5*-specific sgRNA sequences and overhangs for BpiI cut ligation into shuttle vectors pDGE332, pDGE333, pDGE335, and pDGE337 were ordered from Merck (Darmstadt, Germany). The assembled shuttle vectors were handled according to the polyclonal approach recommended in the provided instructions (Stuttmann et al., 2021) and used for BsaI cut ligation into the binary vector pDGE347. To confirm the correct sgRNA sequences, the final vector was sequenced with the recommended primers and transformed into *Agrobacterium tumefacie*ns for *Arabidopsis* floral dip transformation (Clough and Bent, 1998). Transgenic T_1_ seeds were selected using a stereoscope with RFP filter. Transgenic T_1_ plants were genotyped with primers for deletion of the *PML5* locus (FEK_1215/FEK_1218), positive PCR products were Sanger sequenced (Eurofins, Ebersberg, Germany) after gel extraction of the band. In T_2_, non-red seeds were selected, and the plants grown were genotyped again for *pml5* deletion and absence of the pDGE347 transgene (FEK_989/FEK_1095). In T_3_ and T_4_ generations, genotyping was repeated to confirm the results.

### T-DNA lines

SALK_0691926 T-DNA line for *At1g61300*/*PML5* was obtained from the *Nottingham Arabidopsis seeds centre* (*NASC*). The insertion site was confirmed by sanger sequencing and is at 2289bp of the CDS.

### RT-PCR

50µg of 6-week-old *A. thaliana* leaf tissue were collected in a 2-mL reaction tube, frozen in liquid N_2_ and crushed in a bead mill using a stainless-steel bead. Total RNA was extracted using the RNeasy kit (Qiagen). RNA concentration was determined, and equal amounts were used to perform a DNAse I (Thermo Scientific) digest for 30min at 37°C. Again, RNA concentration was determined, and equal amounts were used for RT-PCR with Superscript II reverse transcriptase (Thermo Fisher Scientific; Waltham, USA) according to the manufacturer’s protocol. 2µL of RNA or cDNA were used for PCRs. In addition, 2µL of genomic DNA extracted with SENT buffer (Edwards et al., 1991) was used as a control experiment.

### Biomass quantification

Eight-week-old Arabidopsis plants grown in short days were used. The above-ground tissue was cut and weighed. Graphs were plotted using GraphPad Prism 9.

#### Transient expression in *N. benthamiana*

*Agrobacterium tumefaciens* overnight cultures were centrifuged and resuspended in induction buffer (10mM MgCl_2,_ 10mM MES pH5.6, 150µm acetosyringone). The optical density at 600nm (OD_600_) of all constructs was adjusted to 0.3 and for *35S::P19* to 0.05. *35S::P19* was mixed with all the constructs to be infiltrated, to avoid any possible silencing issues. *Agrobacterium* mixtures were hand-infiltrated into the abaxial site of leaves of 4-5 weeks old *N. benthamiana* plants. *Citrine-HA* or *35S::GFP* single infiltrations were always done as control infiltrations for the experiments. Protein expression was induced at 20 hours post-infiltration by spraying using indicated estradiol (Sigma-Aldrich) concentration and 0.001% (v/v) Silwet L-77.

### Cell death assay

*Nicotiana benthamiana* leaves infiltrated with *Agrobacterium tumefaciens* cultures containing the specific constructs were imaged for cell death at indicated time points post estradiol induction. Cell death images were taken under UV light using an Amersham ImageQuant 800 and an integrated Cy5 filter (GE Healthcare; Chalfont St. Giles, UK).

Arabidopsis seedlings were sown on ½ MS agar plates supplemented with 20µM estradiol and grown for 10 days at long-day conditions (16-hour light/8-hour dark in 21°C/18°C, 45% humidity). Growth defects were photographed using a Canon EOS 80D DSLR camera.

### Immunoblotting

For total protein extraction, leaf tissue from *N. benthamiana* or Arabidopsis was ground using a tissue lyzer (Mill Retsch MM400, Retsch GmbH, Haan, Germany) and resuspended in 190 µl ice-cold grinding buffer (20 mM Tris-HCl pH-7, 150 mM NaCl, 1 mM EDTA pH-8, 1% (v/v) Triton X-100, 0.1% (w/v) SDS, 5 mM DTT, 1X Halt^TM^ PIC (Thermo Fisher Scientific; Waltham, USA). Samples were incubated on ice for 10min and then centrifuged for 15 min at 16000 *g* and 4^0^C. Then 5X SDS loading buffer (250 mM Tris- HCl pH 6.8. 50% (v/v) glycerol, 500 mM DTT, 10% (w/v) SDS, 0.005% (w/v) bromophenol blue) was added to the supernatant. Proteins were denatured at 95°C for 5 min. Proteins were separated into 8% or 10% sodium dodecyl sulfate (SDS) polyacrylamide gels. The proteins were transferred to Amersham^TM^ Protran^TM^ 0.45um Nitrocellulose 300 mm membranes for western blotting. Primary and secondary antibody dilutions used for immune detection were as follows: α-HA(rat)-1:2,000 (Roche Diagnostics, Basel, Switzerland), α-GFP(mouse)-1:1500 (Roche Diagnostics, Basel, Switzerland), α-RFP(rat)-1:1,000 (ChromoTek, Planegg-Martinsried, Germany), α-Myc(rat)-1:1,000 (ChromoTek, Planegg-Martinsried, Germany), α- HA(rat)-1:2000 (Roche Diagnostics, Basel, Switzerland), α-UGPase(rabbit)-1:2,000 (Agrisera, Vännäs, Sweden), α-Histone H3(rabbit)-1:5,000 (Agrisera, Vännäs, Sweden), α-mouse HRP-conjugated 1:10,000 (Sigma-Aldrich, St. Louis, USA), α-rat HRP-conjugated 1:10,000 (Thermo Fisher Scientific; Waltham, USA), α-rabbit HRP- conjugated 1:10,000 (Sigma-Aldrich; St. Louis, USA). Chemiluminescence was detected using an Amersham ImageQuant 800 (GE Healthcare; Chalfont St. Giles, UK). Images were processed with Coral Photo Paint (Corel Corporation; Ottawa, Canada) to adjust brightness and contrast.

### Co-immunoprecipitation

*Nicotiana benthamiana* leaf samples were frozen and ground in liquid nitrogen. Tissue powder was resuspended in 2.5 ml extraction buffer (50 mM HEPES buffer pH 7.5, 50 mM NaCl, 10 mM EDTA pH 8.0, 0.5% [v/v] Triton X-100, 5 mM DTT, 1x Halt^TM^ Protease Inhibitor Cocktail (Thermo Fisher Scientific; Waltham, USA)). Samples were kept on ice for 20 min and centrifuged at 16,000 g for 15 min. The supernatant was subjected to immunoprecipitation for 1 hour using anti-GFP (ChromoTek, Planegg- Martinsried Germany), anti-RFP (ChromoTek, Planegg-Martinsried Germany) or anti- Myc beads (ChromoTek, Planegg-Martinsried Germany). Beads were spun down by centrifugation at 2,400 g at 4°C and washed two times with 1ml wash buffer (50 mM HEPES buffer pH-7.5, 150 mM NaCl, 10 mM EDTA pH-8.0, 0.2% [v/v] Triton X-100, 5 mM DTT, 1x Halt^TM^ Protease Inhibitor Cocktail (Thermo Fisher Scientific; Waltham, USA)) by incubating the extracts for 5min on a rotating wheel at 4°C and two additional times by inverting the tube six times. Bound proteins were eluted in 120μl 2x SDS loading buffer (100 mM Tris-HCl pH-6.8, 20% [v/v] glycerol, 200 mM DTT, 4% [w/v] SDS, 0.002% [w/v] bromophenol blue) and denatured by boiling the proteins at 95°C for 5 min.

### Microsomal fractionation

Microsomal membrane fractions were prepared from 100-200 mg *N. benthamiana* leave tissue transiently expressing the indicated construct. Plant tissue as ground in liquid nitrogen and 2 ml sucrose buffer (20 mM Tris pH 8.0, 0.33 M sucrose, 1 mM EDTA, 5 mM DTT and 1X Halt^TM^ PIC (Thermo Fisher Scientific) was added. Samples were centrifuged at 2,000 *g* for 10 min and the supernatant were used for the following fractionation. 120 µl were taken away and used as the total protein extract sample (T) and the remaining supernatant was subjected to centrifugation at 17,000 g for 1 hr at 4°C. The pellet obtained was resuspended in 50 µl sucrose buffer and represents the microsomal/membrane fraction (M). The supernatant is the cytosolic fraction (S). The fractions obtained were used for SDS-PAGE.

### Blue Native PAGE

*Nicotiana benthamiana* leaf discs (n=∼13 (4 mg per disc), 50 mg) were ground in microcentrifuge tubes using a tissue-lyzer (Mill Retsch MM400, Retsch GmbH, Haan, Germany) and mixed with ice cold modified GTEN-DDM buffer (3.6 µl/mg) (10% glycerol,100 mM Tris-HCl, pH - 7.5, 1 mM EDTA, 150 mM NaCl, 5 mM dithiothreitol (DTT), 1X plant protease inhibitor cocktail (Halt^TM^ PIC (Thermo Fisher Scientific), 0.5% (w/v) DDM (n-dodecyl β-D-maltoside) (Avanti polar lipids, Inc, USA)). The mixture was vortexed and placed on ice, then centrifuged at 20,000g in 4°C. The supernatant is the BN-PAGE sample. This supernatant was mixed with 4X native PAGE sample buffer (Invitrogen, Thermo Fisher Scientific, Carlsbad, CA) in the ratio of 3:1. Further add 0.5 µl of NativePAGE 5% G250 sample additive (Invitrogen, Thermo Fisher Scientific, Carlsbad, CA) to the 19.5 µL of the previous mixture and mix it well. This processed sample was used for running the BN-PAGE gels. Immunoblotting and detection of the complexes were carried out using the methods described in (Jacob et al., 2021).

### Confocal Laser Scanning Microscopy

Confocal laser scanning microscopy was performed using a Zeiss (Oberkochen, Germany) LSM 880 laser-scanning confocal microscope equipped with a 40x water immersion objective. Images were obtained and analyzed using Zeiss ZEN blue software for adjusting the brightness and contrast. Citrine and YFP fluorescence were excited at 514 nm, GFP at 488 nm, and RFP at 561 nm and emission was detected at 510-570 nm, 516/519-550 nm, and 588-651 nm, respectively. Chlorophyll A was excited at 561 nm and detected at 656-676 nm. The constructs for the cellular markers used in the study are *p35S::CD3-967-mCHERRY* as Golgi marker (Nelson et al., 2007), *p35S::sp-RFP-AFNY-STOP-tRFP* as a vacuolar lumen marker, *pUBN::RFP* as a cytosolic marker, and *p35S::BRI1-RFP* as plasma membrane marker.

### Bimolecular fluorescence complementation (BiFC)

For BiFC, *Agrobacterium tumefaciens* (OD_600_ = 0.2) containing the indicated constructs were mixed in a 1:1 ratio, with P19 expressing *A. tumefaciens* (OD_600_ = 0.05), and hand-infiltrated into leaves of 4-week-old *N. benthamiana*. YFP fluorescence was imaged at indicated time points using confocal laser scanning microscopy.

### Bacterial infection assay

*Pseudomonas syringae* strains used in the study were grown on KB agarose plates with relevant antibiotics. Over-night grown bacteria were re-suspended in 10mM MgCl_2_ and diluted to the required final concentration. Bacterial culture was hand infiltrated into rosette leaves of six-week-old plants grown under short-day conditions. Leaf disc samples were collected on day 0 (0 dpi) and day 3 (3 dpi), ground in 200 µl distilled water using a tissue lyzer (Mill Retsch MM400, Retsch GmbH, Haan, Germany). Dilution series of the bacterial culture were plated on KB agarose plates with appropriate antibiotics and cycloheximide, and colonies were counted 2 days later after incubation at 28°C. Replicates were analyzed by one-way ANOVA with post-hoc Tukey’s test using GraphPad Prism 9. Bacterial cultures for cell death assays were hand-infiltrated on the right side of the leaf. Leaves were removed at the indicated time points, and autofluorescence was measured using a Typhoon FLA9500 laser scanner (Cytiva, Chicago, IL) of the adaxial side of the leaf. Image processing was done using ImageJ.

### Virus infection assay

Tobacco rattle virus (Liu et al., 2002) was used. *Agrobacterium* containing pYL156- TRV1 or pYL156-TRV2-EV plasmids were cultured in LB medium (containing 100 mg/L Kan and 25 mg/L Rif) for 18 h at 28°C by shaking at 220 rpm. *Agrobacterium* cells were then collected by centrifugation at 4000 *g* for 15min, followed by resuspension in resuspension buffer (10 mM MgCl_2_, 100 μM acetosyringone, and 10 mM MES, pH 5.8) and incubated at 28°C for 3 h in the dark before infiltration into *N. benthamiana* plants. After 2 weeks, virus-infected leaves were harvested and ground on ice with extraction buffer (10 mM Na2HPO4 / NaH_2_PO_4_, pH 7.2). The extracts were filtered through Miracloth and centrifuged at 4°C for 1min at 12,000 g. Subsequently, the 4-week-old Arabidopsis plants were inoculated directly by rubbing the fresh supernatant of extracts generated from 2-week-old TRV-infected *N. benthamiana* plants onto rosette leaves. The Arabidopsis leaves were harvested at 6 and 8 days after inoculation and analyzed by qRT-PCR using Primer pairs (TRV-F: GTGCACGCAACAGTTCTAATCG and TRV-R: GCTGTGCTTTGATTTCTCCACC; Actin-F: CAGTGTCTGGATCGGTGGTT and Actin-R: TGAACGATTCCTGGACCTGC). The experiments were repeated three times with the similar results.

### PTI experiments (ROS and ethylene measurements)

Leaves of 5-6-week-old Arabidopsis were cut into pieces of equal size and floated on H_2_O overnight. For ROS burst, the leaf pieces were transferred to a 96-well plate (1 piece/well) containing 100 µl of solution with 20 µM L-012 (Waco) and 2 µg ml-1 peroxidase. Luminescence after treatment with peptides or water (as control) was measured with a luminometer (Mithras LB 940, Berthold) in 2 min intervals. Total relative light unit (RLU) production was determined by calculating the area under the scatter curve for the time points indicated. The RLU values at time point 0 min were set as 0 to accordingly remove the background responses. For ethylene production, three leaf pieces were incubated in a sealed 6.5 ml glass tube with 0.4 ml 20 mM MES buffer, pH 5.7 and the indicated elicitors. Ethylene production was measured by gas chromatographic analysis (GC-14A; Shimadzu) of 1 ml air from the closed tube after incubation for 4 hrs.

### In-Silico Analysis

Phylogenetic trees and alignments in Supplementary figures S1, S2, S4 and S5 were constructed using CLC Main Workbench 23.0.1 (Qiagen, Venlo, Netherlands). The structure prediction for PML5 was done using AlphaFold 2 (https://alphafold.ebi.ac.uk/) and the UniProt ID of PML5 is O64790 DRL17_ARATH.

To examine whether the PML5 deletion in Arabidopsis is unique in the Brassicales we constructed a phylogeny using PML5 and homologs without the deletion from other representative Brassicalee genomes. Using Orthofinder (Emms and Kelly, 2019) we clustered the predicted protein sequences from three Capsella genomes (*Capsella bursa pastoris*, *Capsella rubella*, and *Capsella orientalis*, unpublished) and 18 Arabidopsis genomes. Using mafft (Katoh and Standley, 2013) we then aligned the resulting orthogroup containing the PML5 homologs with additional homologs from existing proteomes (*Brassica napus* (BnaCnng57760D), *Capsella rubella* (XP_006300767.1, Carub.0002s0399), *Capsella bursa pastoris* (Cbp33710), *Cardamine hirsute* (CARHR052110), *Leavenworthia alabamica* (LA_scaffold903_5) . We then trimmed the amino acid alignment to just the NBARC region, which does not contain the deletion, and constructed a maximum-likelihood phylogeny using model selection in IQtree that selected the JTT model (model JTT, ultra-fast bootstraps: 1000) (Nguyen et al., 2015).

## SUPPLEMENTAL INFORMATION

Supplemental information can be found online.

## AUTHOR CONTRIBUTIONS

Conceptualization, F.E.K., S.S., S.B., D.W., R.L.D., T.N.; Methodology, S.S., S.B., E.S., K.F., F.W., A.M. C.P., L.T., Z.J., L.Z., E.A.P.; Investigation, F.E.K., S.S., S.B., L.T., Z.J., L.Z., E.A.P..; Writing – Original Draft, S.S. and F.E.K..; Writing – Review & Editing, F.E.K., S.S., S.B., D.W., R.L.D., T.N.; Funding Acquisition, F.E.K., R.L.D., T.N. and D.W.; Resources, F.E.K., R.L.D., T.N. and D.W.; Supervision, F.E.K.

## Supporting information

Supplemental Figures_reduced

## ACKNOWLEDGEMENTS

We thank Bettina Hause for the CD3-967 construct, Manoj K Singh (Gerd Jürgens Lab) for the (UBQ)-pXVE-GFP empty vector and Klaus Harter for the BRI1 construct, Elke Sauberzweig, Christel Kulibaba-Mattern and Patrick Vetter for technical support, the ZMBP gardeners and microscopy facility for their support and advice, Xander Zuijdgeest, Frank Vogt and Svenja Saile for critical reading of the manuscript, and other members of the El Kasmi, Nishimura and Dangl labs as well as Sandra Richter and Paul Gouguet for critical comments and discussions. We appreciate the support of Prabha Manishankar (Oecking Lab) and Nak Hyun Kim (Dangl Lab) for helping us to set up the Blue Native PAGE assay.

## SUPPLEMENTAL FIGURE LEGENDS

Figure S1: Most Arabidopsis ANLs are also PML NLRs

Arabidopsis Col-0 ANLs can be grouped into 5 subclusters, and most ANLs have a putative N-myristoylation and/or S-acylation site at their N-termini (ANLs lacking this site(s) are indicated by a black star). PML5/At1g61300 subclusters in group 3 with PML11/At1g61180, PML12/At1g61310 and PML13/At1g61190. Phylogenetic tree is based on an alignment build with full length protein sequences and was generated with CLC Main Workbench tool v.22.02.

Figure S2: Amino acid alignment of full-length PML5 and its homologs in Col-0

Alignment of PML5/At1g61300 and its closest Col-0 homologs PML11/At1g61180, PML12/At1g61310 and PML13 At1g61190. Differences in amino acid composition are highlighted in light green and amino acid conversation (in percentage) is presented as a bar-blot in blue. Alignment was done with CLC Main Workbench tool v.22.02.

Figure S3: PML5 functions as a canonical cell death inducing NLR.

(A) AlphaFold 2 structural prediction of PML5. CC, NB-ARC, and LRR domains are indicated. Potential alpha-helix is predicted between Asp/D13 and Ile/I 27. Colors indicate confidence; red = very low, yellow = low, light blue = confident and dark blue = very high. (B) Sequence alignment of the first 26 N-terminal amino acids of PML5 with NRC4(Nb) and ZAR1(Col-0), highlighting the hydrophobic amino-acids (C) Cell death induced by transiently expressed wildtype PML5 and mutant variants constitutively expressed under 35S promoter. a Protein expression analysis (lower panel) of constructs infiltrated in (C) Citrine-HA was used as a negative control. Pictures were taken 2 days post-infiltration. (D) Cell death phenotype by transiently expressed wildtype PML5 and mutant variants conditionally expressed under a 35S promoter-controlled estradiol inducible system. Protein expression analysis of indicated constructs is shown in the lower panel. Citrine-HA was used as a negative control. Pictures were taken 3 days post-induction (dpi). (C) and (D) Total protein extracts were immunoblotted and detected with an anti-GFP (α-GFP) antibody. The lower blot shows singular eYFP-HA (about 30 kDa). Ponceau S (PS) staining of the Rubisco protein band is shown as a loading control. In (C), WT = wildtype PML5; D362V = MHD mutant; K74R = P-loop mutant; GCC(2,3,4)A = S-acylation mutant; GCC/2,3,4)A,D362V = S-acylation and MHD mutant. (E) Cell death phenotype (in false color) in a single rosette leaf of four-week-old Arabidopsis plants at 18hpi with 20µM estradiol is shown. Red indicates strong cell death in PML5 WT and D362V lines and green shows no cell death in K74R and GFP negative control lines. (F) Growth restriction/Cell death phenotype of two independent transgenic *Arabidopsis* lines conditionally overexpressing PML5-GFP, D362V-GFP. A *35::GFP* plant line was used as a negative control. 10-day-old Arabidopsis seedlings grown on ½ MS plates supplemented with or without 20µM estradiol are shown.

Figure S4: Alignment of PML5 and PML5-like NLRs of different Arabidopsis accessions.

Alignment of PML5/At1g61300 and PML5-like NLRs of 13 Arabidopsis accessions. The close Col-0 homolog PML11/At1g61180 is shown to better visualize the 113 amino acid deletion. Differences in amino acid composition are highlighted in light green and amino acid conversation (in percentage) is presented as a bar-blot in blue. Alignment was done with CLC Main Workbench tool v.22.02.

Figure S5: PML5-like deletion may have occurred independently in

***Arabidopsis* and *Capsella* species.**

A maximum likelihood phylogeny based on the amino acid sequence of the NBARC domain of PML homologs from representatives of the Brassicales. Nodes with black dots have a bootstrap support of at least 85%. The multiple sequence alignment next to the phylogeny shows the presence and absence of the 5-prime deletion across the tree. Arabidopsis PML5 sequences are nested within the intact homologs present in the species.

Figure S6: PML5-induced cell death is independent of RNL helper NLRs and EDS1.

(A) Cell death induced by transiently expressed wildtype (WT) and PML5 D362V mutant in WT, *eds1, adr1, nrg1, adr1 nrg1 N. benthamiana* mutant lines. The RPM1 D505V auto-active mutant was used as a positive control and Citrine-HA as a negative control. Images were taken 1-day post-induction with 20µM estradiol. Leaves are shown in false color, red indicates cell death and green healthy/alive tissue.

Figure S7: PML5 N-terminal 60 amino acids are sufficient for cell death induction.

(A) Cell death induced by transiently expressed full-length PML5 and PML5 domains in *N. benthamiana* leaves. Black/Grey coloring indicates cell death. Note: only full- length WT PML5 and its CC^1-60^ domain were inducing cell death symptoms. eYFP-HA was used as a negative control. Image was taken 3 days post-induction with 20µM estradiol. (B) Combination of PML5 CC, NB and LRR domains is not sufficient to induce a WT-like cell death response. PML5 domains were transiently expressed alone or in co-infiltrations *in N. benthamiana.* Image was taken 2 days post-induction with 20µM estradiol. (Lower panel) Total protein extracts were immunoblotted and detected with an anti-HA (α-HA) antibody. Ponceau S (PS) staining of Rubisco protein band is shown as loading control. The dots indicated are the different fragments expressed, Red = co-infiltrated singular eYFP, Green = NB domain, Orange = LRR domain, and Yellow = CC domain. (C) PML5 CC^1-60^ self-associates in *N. benthamiana* transient expression. C-terminally YFP and MYC tagged CC^1-60^ were co-expressed. Total proteins were immunoprecipitated using anti-GFP (α-GFP) beads and immunoblotted using an anti-GFP (α-GFP) and anti-Myc (α-myc) antibody. Ponceau S (PS) staining of Rubisco protein band is shown as loading control. (D) Protein-blot analysis of total proteins from the transiently expressed proteins of the BiFC assay in main figure 2D. Proteins were detected using anti-Myc (α-MYC) and anti-HA (α-HA) antibodies. Ponceau S (PS) staining of Rubisco protein band is shown as loading control.

Figure S8: PML5 loss of function mutants are mainly cytosolic.

(A-D) Confocal laser scanning microscopy showing subcellular localization of transiently expressed PML5-Citrine-HA mutants in comparison to a nuclear-cytosolic mRFP; K74R (A); GCC(2,3,4)A (B); GCC(2,3,4)A D362V (C) and D362V (D). All loss of function mutants exhibits a cytosolic distribution. The D362V mutant has a similar localization pattern as the PML5 WT, and induces an, even more, severe vacuolar vesiculation phenotype (D). White dotted lines indicate the area used for colocalization profile analysis and the corresponding profiles are shown on the right. n = nucleus. Scale bars, 20uM; for information on compartment marker proteins see the material and method section. Protein expression was induced with 20 uM estradiol and confocal imaging was performed at 8 hours post-induction for K74R; 9hpi for GCC(2,3,4)A; 9hpi for GCC(2,3,4)A, D362V and 4hpi for D362V. (E) Subcellular fractionation of transiently expressed PML5-citrine-HA protein indicates a strong microsomal/membrane association. Total protein extracts (6hpi) were immunoblotted and detected with an anti-HA (α-HA) antibody for PML5, anti-UGPase (α- UGPase) as a cytosolic marker, and anti-Histone H3 (α-Histone H3) as a microsomal marker. T = total protein fraction; S = soluble protein fraction; M = microsomal protein fraction.

Figure S9: *PML5* does not contribute to resistance against avirulent *Pst* DC3000.

Rosette leaves of six-week-old *Arabidopsis* plants were hand infiltrated with various *Pst* DC3000 strains using an OD OD_600_ = 0.001 (A) *Pst* DC3000, (C) *Pst* DC3000 *AvrPphB*, (E) *Pst* DC3000 *HopZ1a*, (G) *Pst* DC3000 *AvrRps4*, (I) *Pst* DC3000 D36E and bacterial growth was determined on day 0 and day 3 post infiltration. Data points (colony forming units per square cm – cfu/cm^2^) are shown as black dots and represent 3 biological replicates with each 4 technical replicates (*n*=12, each with 6 leaf samples). Tukey’s HSD test was performed (α = 0.05). (B, D, F, H) Induction of HR is not affected in *pml5* mutants. The right side of the leaves was hand infiltrated with *Pst* DC3000: *AvrRpm1*(B)*, AvrPphB* (D) and *HopZ1a* (F) at an OD_600_ = 0.1 and with *AvrRps4* (H) at an OD_600_ = 0.2. Infiltrated leaves were imaged with a Typhoon laser scanner at 5 hours post infiltration (hpi) for *AvrRpm1*(B), 22hpi for *AvrPphB (*D), 24hpi for *HopZ1a (*F) and *AvrRps4* (H). The leaf images are shown in false color: Purple/blueish parts indicate non-infiltrated healthy tissue, and orange/yellowish dead cells. (J) Infiltration of *Pst* DC3000 D36E does not induce visible disease symptoms on *pml5-1*, *pml5-2c* or Col-0 and *eds1-12*.

Figure S10: Loss of *PML5* has no effect on PRR-induced responses and TRV resistance.

(A) Total ROS production in leaf discs of Col-0 and *pml5-2c* mutant treated with water (mock), 1µM nlp20, or 100nM flg22 over 60 min. RLU, relative light unit. Data points are indicated as grey dots from four independent experiments (n=28) and plotted as box plots. (B) Ethylene accumulation in Col-0 and *pml5* mutant after 4 h treatment with water (mock), 1µM nlp20, or 1µM flg22. Data points are indicated as grey dots from five independent experiments (n=15) and plotted as box plots. (C) TRV viral accumulation at 6dpi (left) and 8dpi (right) tested on Col-0, *pml5-1c, pml5-2c* and Empty Vector (EV) *Arabidopsis* plants. Data points are indicated as grey dots from 10 different independent lines and this experiment was repeated 3 times.

