## Supplemental Figures_reduced for "An atypical endomembrane localized CNL-type immune receptor with a conserved deletion in the N-terminal signaling domain functions in cell death and immunity"

### Supplemental Figure S1

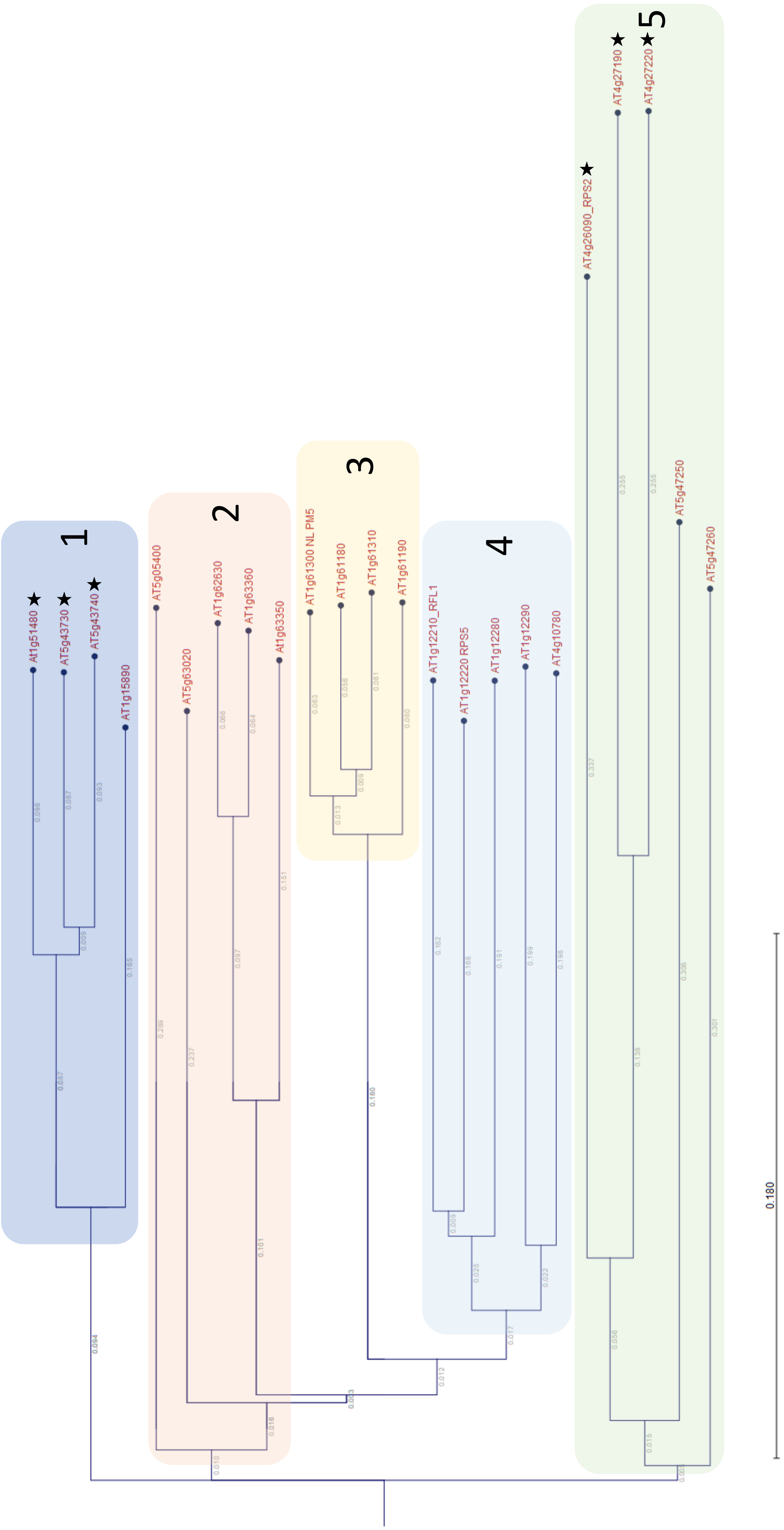

### Supplemental Figure 2

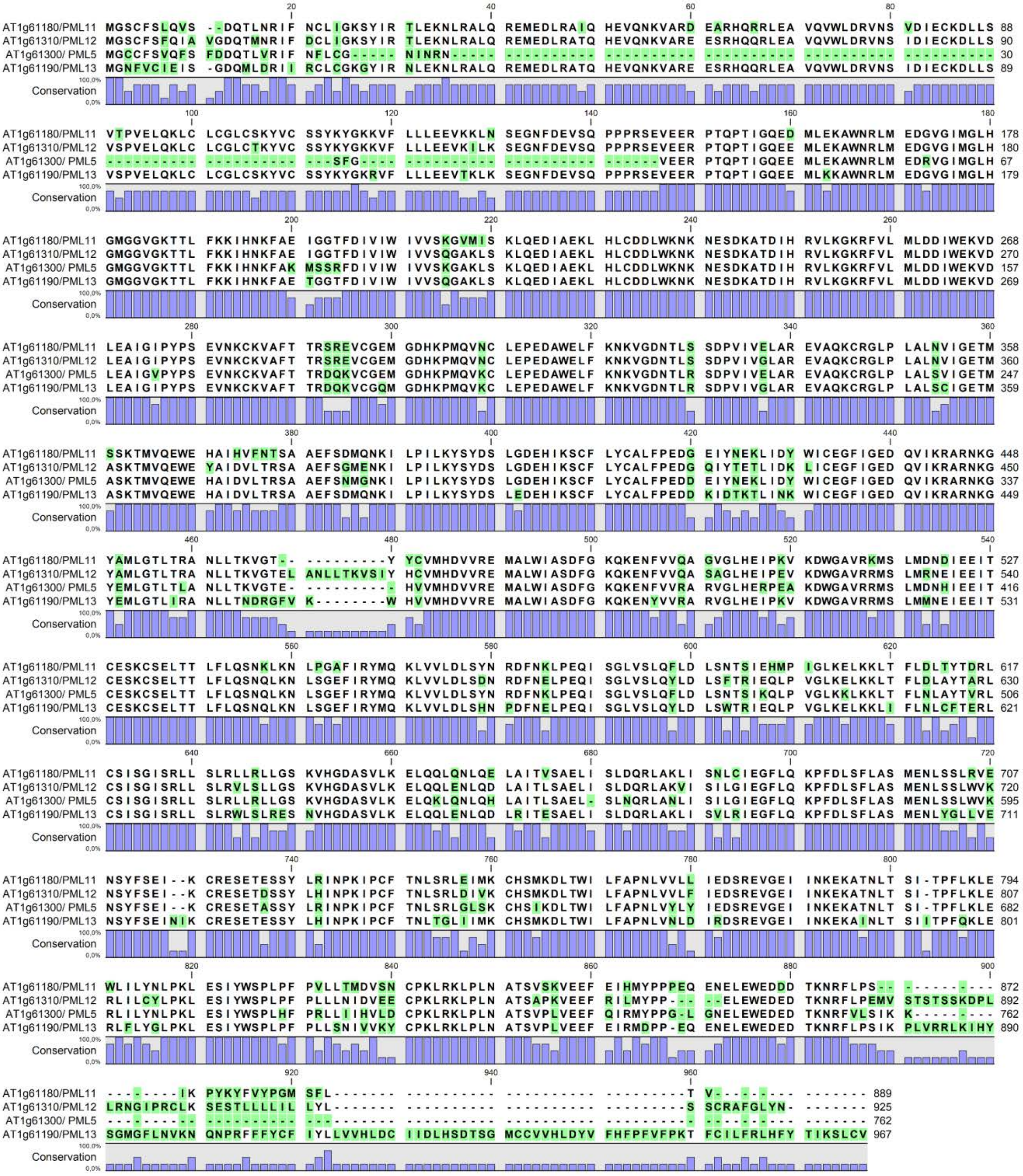

### Supplemental Figure 3

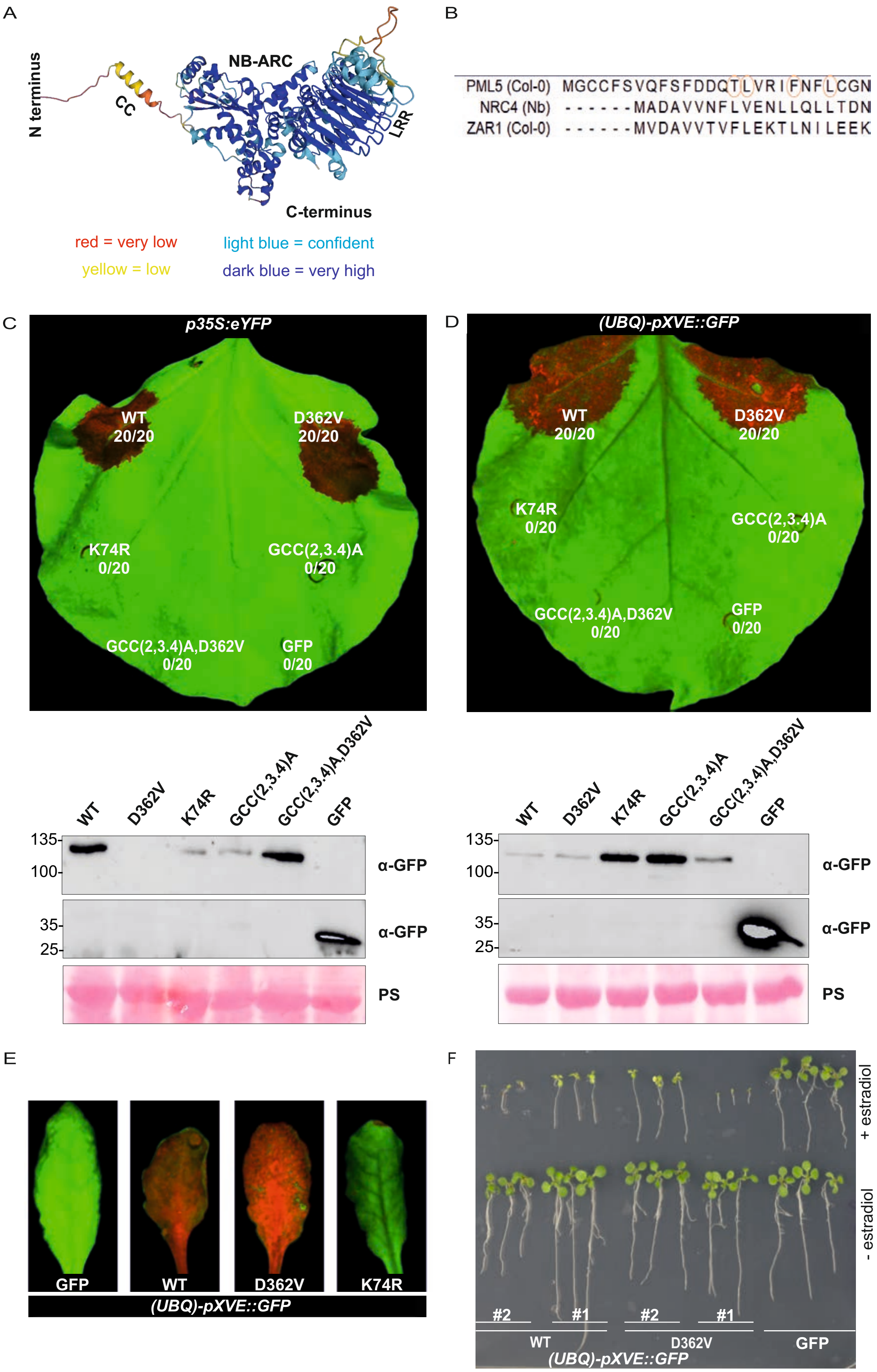

### Supplemental Figure S4

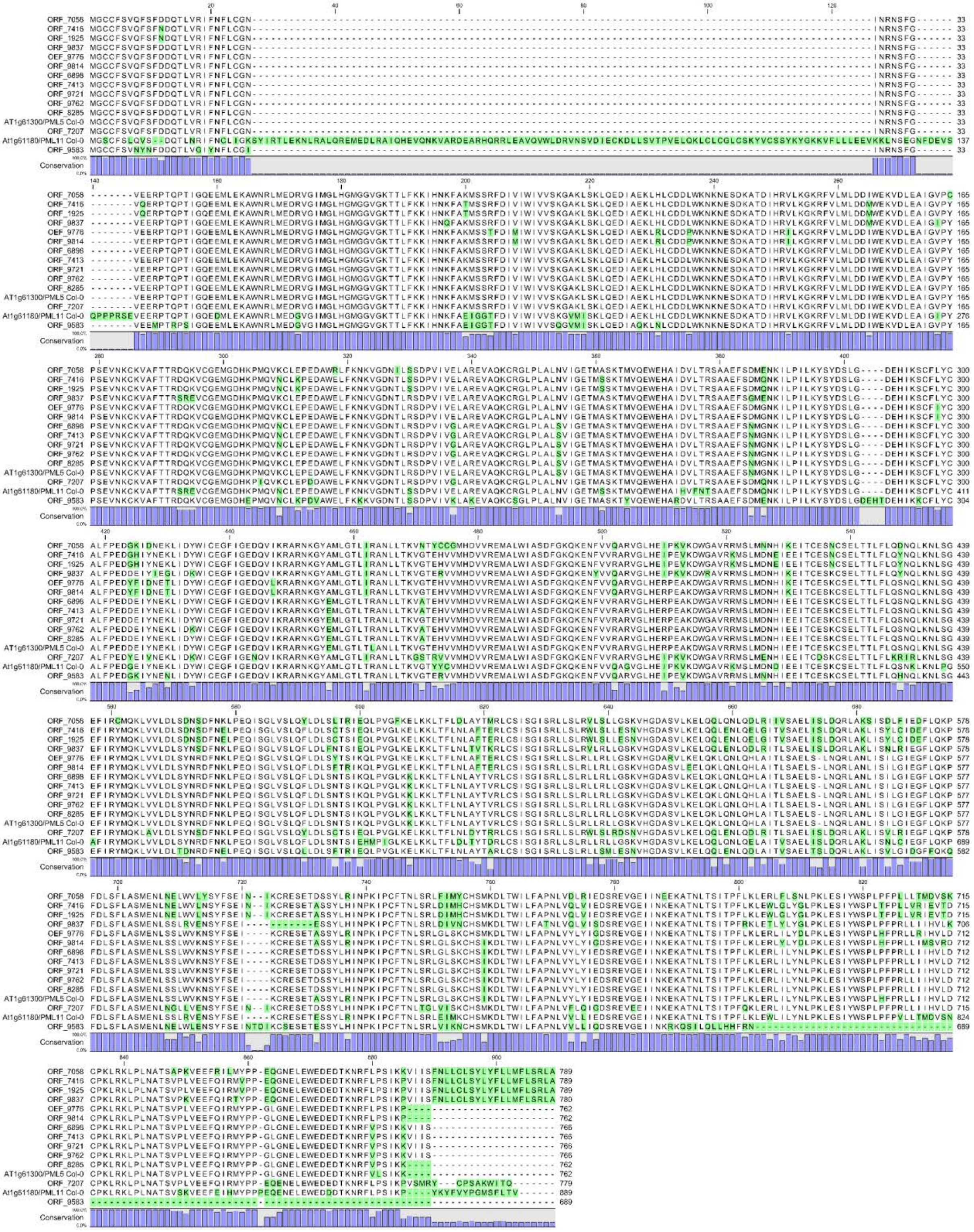

### Supplemental Figure S5

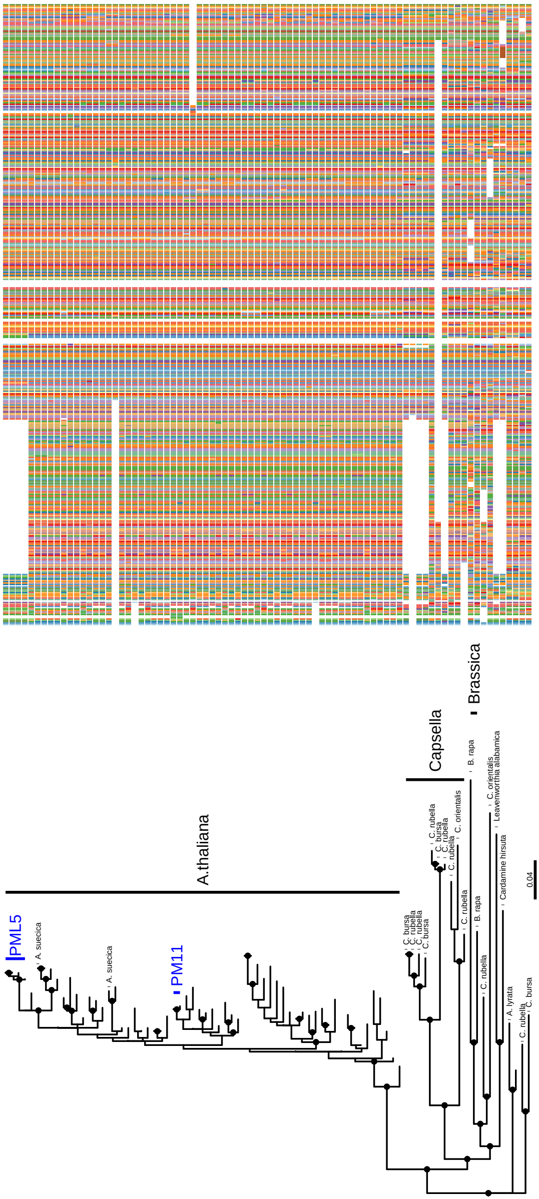

### Supplemental Figure S6

A

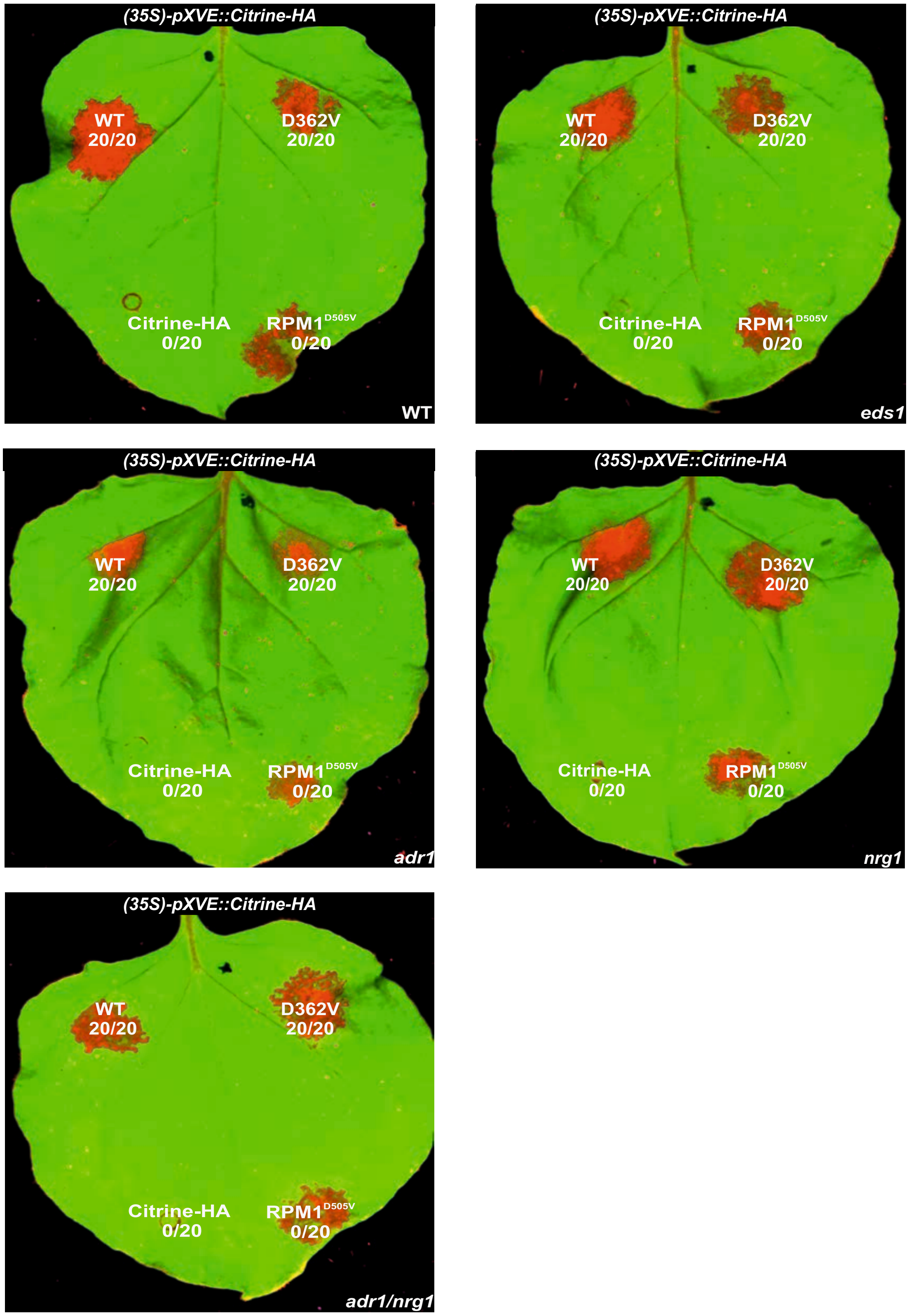

### Supplemental Figure S7

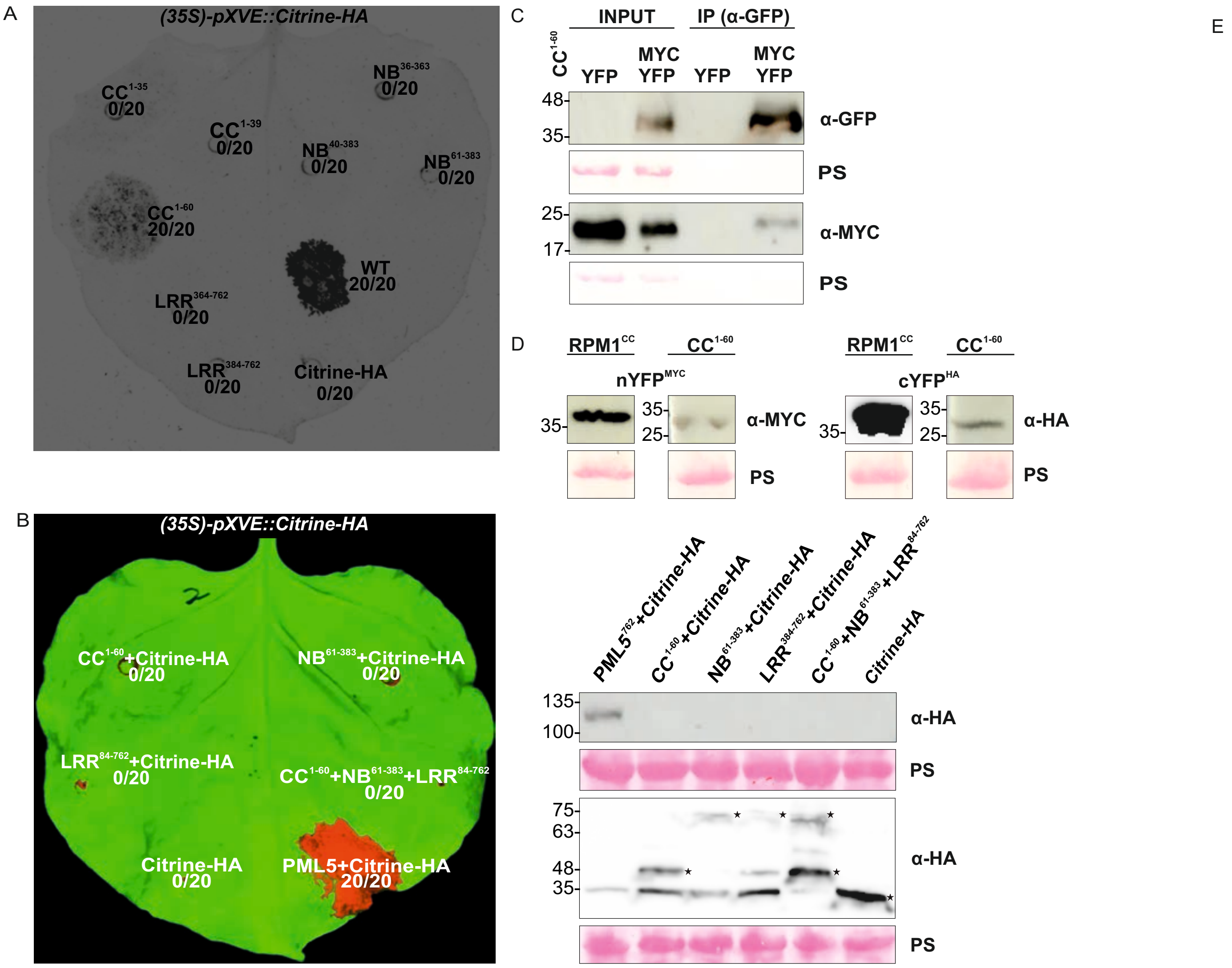

### Supplemental Figure S8

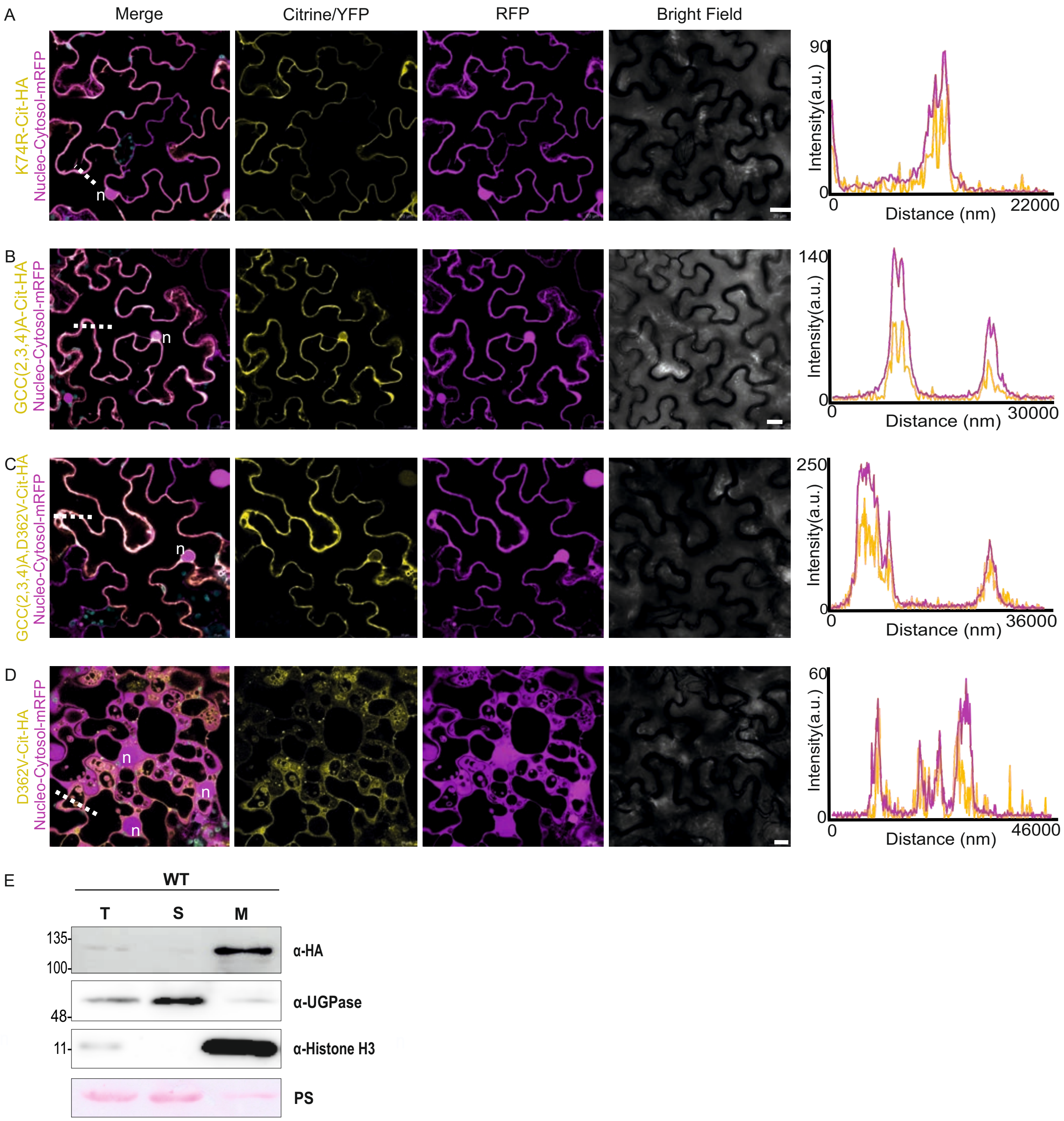

### Supplemental Figure S9

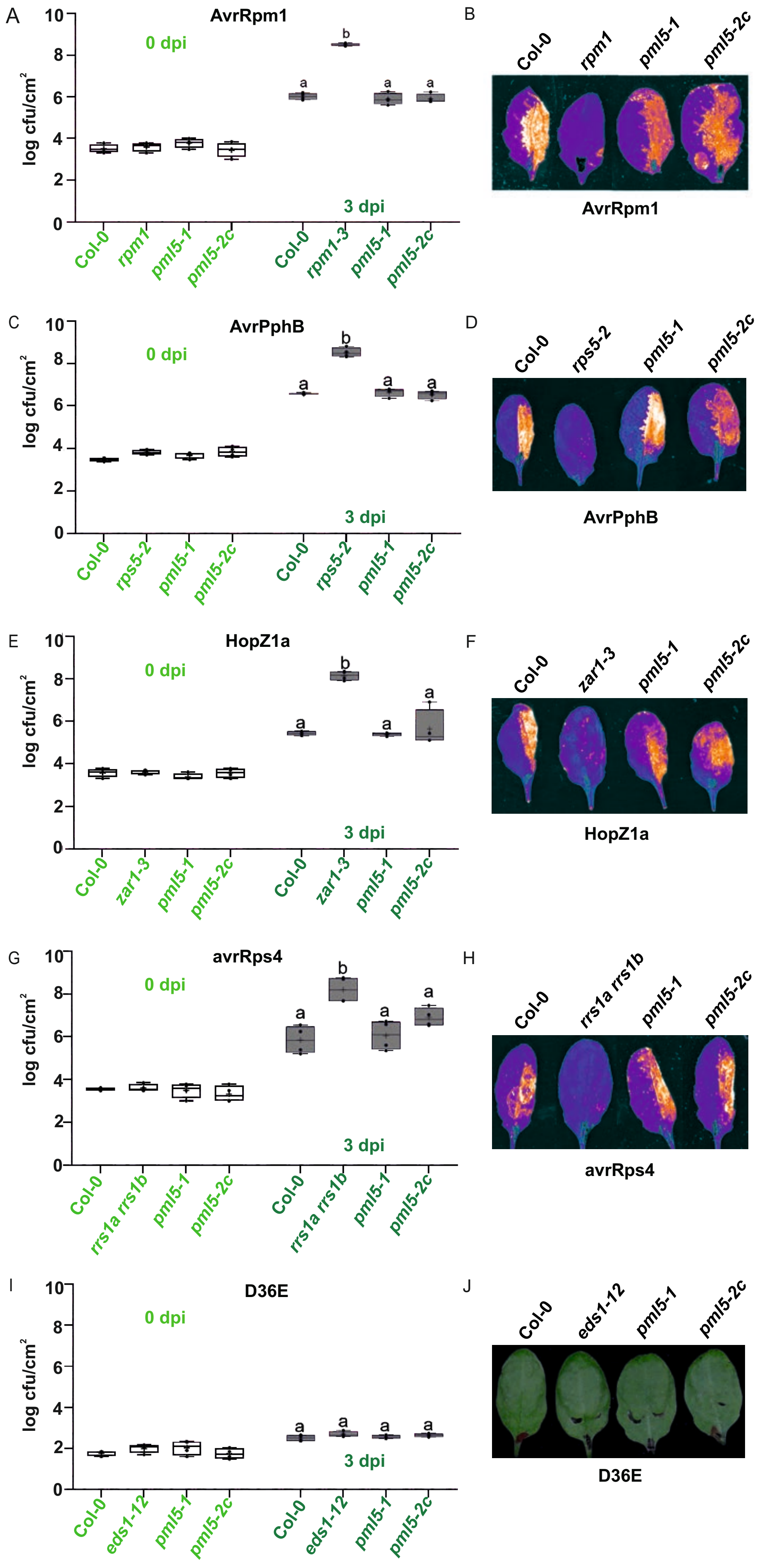

### Supplemental Figure S10

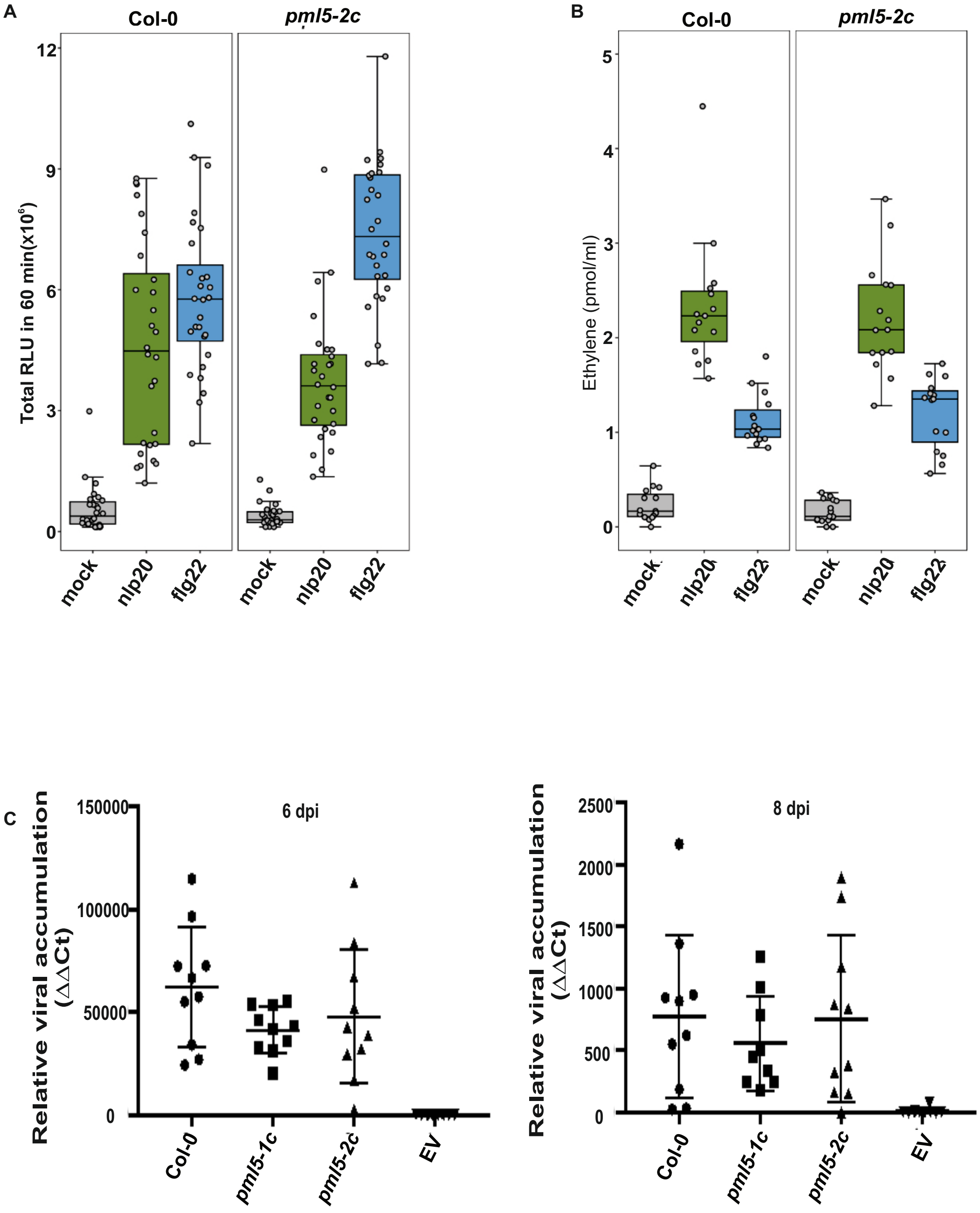
